# Inhibition of host *N*-myristoylation compromises the infectivity of SARS-CoV-2 due to Golgi-bypassing egress from lysosomes and endoplasmic reticulum

**DOI:** 10.1101/2023.03.03.530798

**Authors:** Saber H. Saber, Mohammed R. Shaker, Julian Sng, Nyakuoy Yak, Sean D. Morrison, Selin Pars, Huiwen Zheng, Giovanni Pietrogrande, Tobias Binder, Siyuan Lu, Matthias Floetenmeyer, Ravi Ojha, Tania Quirin, Janika Ruuska, Teemu Smura, Tomas Strandin, Ravi Kant, Lauri Kareinen, Tarja Sironen, Gert Hoy Talbo, Yanshan Zhu, Kirsty R. Short, Jessica Mar, Wouter W. Kallemeijn, Edward W. Tate, Roberto Solari, Ashley J. van Waardenberg, Olli Vapalahti, Ernst Wolvetang, Giuseppe Balistreri, Merja Joensuu

## Abstract

Severe acute respiratory syndrome coronavirus 2 (SARS-CoV-2), which caused the coronavirus disease 2019 (COVID-19) pandemic, remains a global health concern despite vaccines, neutralizing antibodies, and antiviral drugs. Emerging mutations can reduce the effectiveness of these treatments, suggesting that targeting host cell factors may be a valuable alternative. *N*-myristoyltransferases (NMT) are essential enzymes for protein *N*-myristoylation, affecting stability, interaction, localization, and function of numerous proteins. We demonstrate that selective inhibition of host cell NMT decreases SARS-CoV-2 infection by 90% in human lung and primary nasal epithelial cells, and choroid plexus-cortical neuron organoids. NMT inhibition does not affect viral entry, replication or release, but impairs the maturation and incorporation of viral envelope proteins into newly assembled virions, leading to compromised infectivity of released virions. The inhibition of host NMT triggers a Golgi-bypassing pathway for SARS-CoV-2 progeny virion egress, which occurs through endoplasmic reticulum and lysosomal intermediates.

## Introduction

The infection cycle of SARS-CoV-2, the causative agent of the COVID-19 pandemic, starts with the binding of the viral surface trimeric protein spike (S) to the main cell surface receptor angiotensin-converting enzyme 2 (ACE2)^1^. Once attached to the cell, the S must be cleaved by cellular proteases to undergo a conformational change that leads to the fusion of the viral lipid envelope with the membrane of the cell^2^. This fusion event results in the delivery of the viral genome in the cytoplasm of the cell. The cleavage of the viral S is performed by serine proteases such as TMPRSS2, localized at the cell surface or early endosomes^3^, or by cysteine proteases such as Cathepsin-L, localized in the lumen of lysosomes^4^. Upon delivery into the cytoplasm of the target cell, the mRNA-like viral genome is translated by the ribosomes to produce the viral replication machinery^5^. This multiprotein complex, in turn, recruits the newly translated RNA genome to the endoplasmic reticulum (ER) membranes where virus-induced membrane enclosed compartments are then formed^5,6^. Inside these compartments the viral genome is first copied into a complementary strand, which serves as a template for shorter mRNA-like transcripts, the sub-genomic RNAs, as well as for the synthesis of full-length viral genomes that are released into the cytoplasm^5,6^. Among the genes encoded by the sub-genomic mRNAs are the four viral structural proteins, S, Envelope (E) and Membrane (M) protein, which are inserted in the viral envelope membrane, and the nucleoproteins (N) that bind to the newly synthesized viral RNA genomes to form a ribonucleoprotein complex^5^. The final assembly of the viral particle takes place at modified cellular membranes derived from ER, Golgi, and ER–Golgi intermediate compartment (ERGIC)^7^. In ER and Golgi, the S protein undergoes extensive glycosylation which significantly impacts its folding, stability, and interaction with ACE2^8,9^. Additionally, S proteins are further cleaved at a multibasic site into S1 and S2 subunits by the Golgi enzyme Furin, presumably in the trans-Golgi network (TGN)^10^. The virions that initially bud into the lumen of the ERGIC are released from the cells through a process that is not fully characterized and appears to involve transport through lysosomes, before calcium-dependent secretion^11,12^.

*N*-Myristoylation is a co-and post-translational lipid-modification of proteins catalysed by the enzymes *N*-myristoyltransferases (NMT1 and NMT2). This modification plays a crucial role in targeting proteins to cellular membranes, particularly the plasma membrane and membranes of ER and Golgi apparatus^13^. The most common type of *N*-myristoylation occurs at N-terminal glycine of the nascent chain of the protein, emerging from the ribosome,, at a consensus motive (M)GXXXXS_6_ (M is methionine, G glycine, X any amino acid, and S serine)^14^. However, *N*-myristoylation can also occur at an internal amino acid residue when a protein is proteolytically cleaved exposing a G, or potentially a K, at the new N-terminus of one of the cleavage products^14,15^.

In humans, *N*-myristoyltransferases exist in two isoforms, NMT1 and NMT2, which are highly conserved across all mammals ^16,17^. The life cycle of several pathogens, including various bacteria and viruses, depends on protein N-myristoylation^18^. *N*-Myristoylation of viral proteins is observed or predicted in families ranging from RNA to large DNA viruses, including *Picornavirdae, Arenaviridae, Reoviridae, Hepadnaviridae, Polyomaviridae, Ascoviridae, Herpesviridae, Poxviridae, Asfiviridae,* and *Iridoviridae*^18^. In all reported cases, the antiviral effect of NMT inhibitors has been attributed to the prevention of direct *N*-myristoylation of viral proteins^19–21^.

The genome of SARS-CoV-2 does not harbour sequences encoding obvious *N*-myristoylation sites. However, *N*-myristoylation of cellular proteins is important for intracellular trafficking, for instance removal of *N*-myristoylation in ADP-ribosylation factor 1 (ARF1) disassociates the protein from Golgi membranes, thus inhibiting the secretion of cytokines and chemokines^22,23^. We, therefore, hypothesized that inhibiting host NMT could affect many host proteins that could potentially regulate one or more steps of a virus life cycle. Herein, we evaluated the antiviral activity of IMP-1088, a potent and selective inhibitor of cellular *N*-myristoyltransferases NMT1 and NMT2 (NMT)^19,24^. Our findings demonstrate that inhibiting NMT1 and NMT-2 in human lung and primary nasal epithelial cells, as well as in human 3D choroid plexus-cortical neuron organoids, disrupts viral assembly and maturation by interfering with the host cell’s secretory pathway membrane trafficking. We reveal that IMP-1088 treatment causes the dispersion of ERGIC and Golgi structures, blocking the trafficking of the virus from the ER to the Golgi, and triggering an alternative Golgi-bypass pathway for SARS-CoV-2 progeny virion egress, occurring through a lysosomal intermediate. Consequently, SARS-CoV-2 progeny virions released from IMP-1088-treated cells show compromised infectivity, containing less S protein, which is also less cleaved by Furin. We further demonstrate the antiviral activity of IMP-1088 against SARS-CoV-2 Omicron and Delta variants, and respiratory syncytial virus (RSV). Together, our findings describe an unconventional Golgi-bypass pathway as the secretion mechanism of defective viral particles in NMTinhibited human cells and reveal the antiviral potential of IMP-1088 in combating SARS-CoV-2 and RSV infections by targeting host cell functions without inducing cytotoxicity.

## Results

### Inhibition of host NMTs prevents spreading of SARS-CoV-2 infection in human lung cells

We began by investigating the potential antiviral effect of IMP-1088 in human lung small carcinoma cells that were engineered to stably express the virus receptor ACE2 and protease TMPRSS2, which cleaves the viral S protein and primes it for fusion (here named A549-AT cells). A549-AT cells were treated with either DMSO (vehicle control) or a range concentrations of IMP-1088 (62.5 nM to 1 µM) at 1.5 hours before (hbi) or 1.5 h post infection (hpi), and infected with a clinical isolate of SARS-CoV-2 corresponding to the ancestral Wuhan (Wuh) strain at multiplicity of infection (MOI) of 0.2 i.u./cell for 24 h. Cells were then fixed and immunostained against viral N protein or dsRNA, the hallmark of viral replication (Fig. 1A). Quantitative analysis using high-content imaging and automated image analysis revealed that nanomolar concentrations of IMP-1088 reduced SARS-CoV-2 Wuh strain infection by approximately 50% compared to the DMSO control at 24 hpi (Fig. 1B, C). A comparable reduction was observed whether IMP-1088 was added 1.5 hours before infection, or 1.5 h after infection, which is a timeframe sufficient for virus entry and delivery of viral genomes into the cytoplasm of target cells^3^.

**Figure 1.**
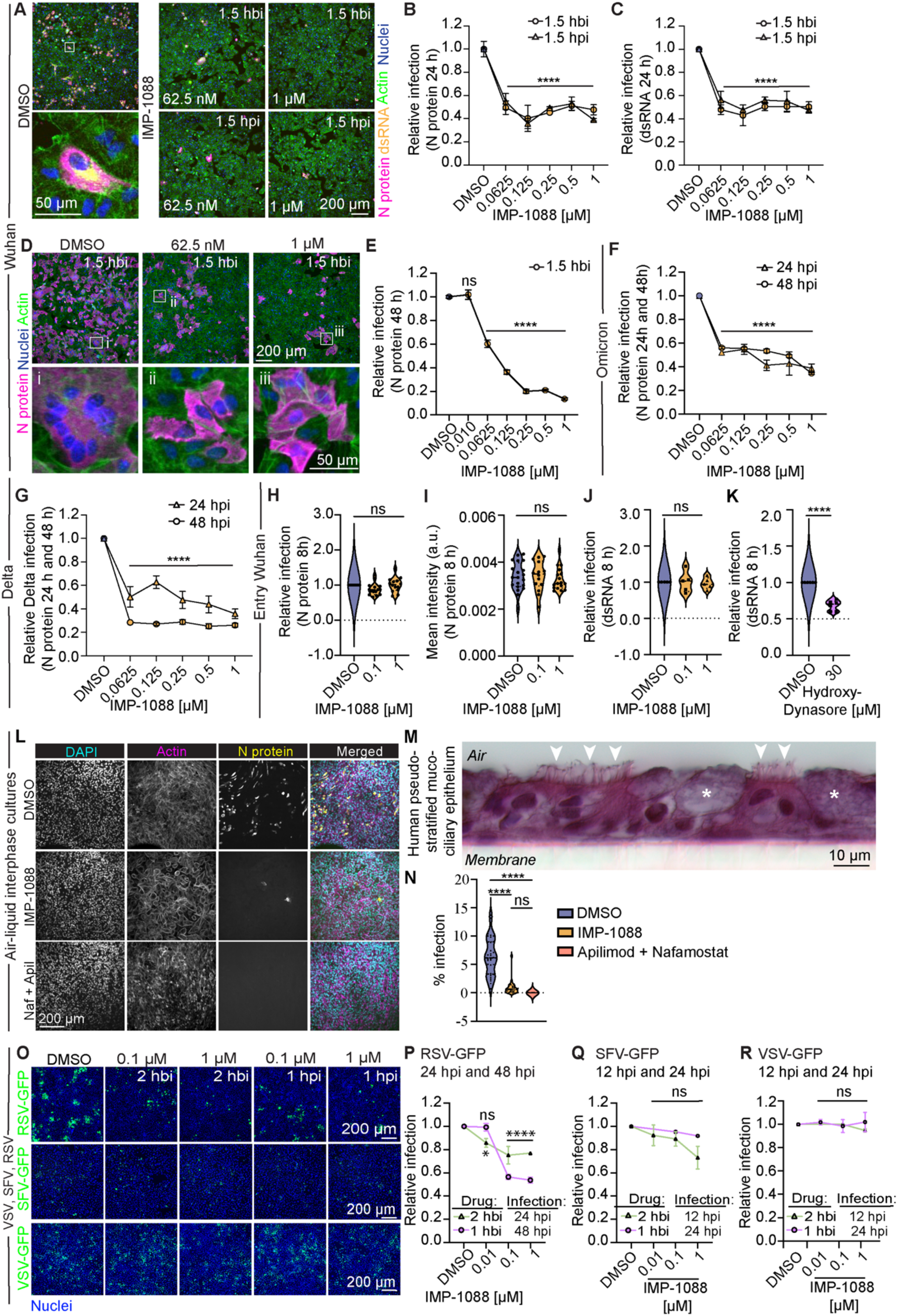
IMP-1088 inhibits SARS-CoV-2 infection in human lung and primary nasal epithelial cells. (A) Representative immunofluorescence staining images of A549-AT lung cells treated with indicated concentrations of IMP-1088 or DMSO vehicle control, 1.5 hours before-infection (hbi) or post-infection (hpi) and infected with SARS-CoV-2 wuh for 24 h. Viral markers N (magenta) and dsRNA (yellow), actin (green) and nuclei (DAPI, blue). (B, C) Quantification of the relative infection (the number of N-positive and dsRNA-positive cells normalized to number of nuclei) in DMSO-and IMP-1088-treated A549-AT cells at indicated concentrations at 1.5 hbi and 1.5 hpi. (D) Representative immunofluorescence images and (E) quantification of SARS-CoV-2 wuh infection (normalized to number of nuclei) in A549-AT cells treated with indicated concentrations of IMP-1088 or DMSO vehicle control 1.5 hbi, infected with for 48 h and stained against N (magenta), actin (green), and nuclei (blue). The boxed areas i (multi-nucleated syncytium of cells), ii and iii (infected single cells) are shown magnified at the bottom of each image. (F, G) Quantification of the relative SARS-CoV-2 Omicron (F) and Delta (G) infection at 24 and 48 hpi in A549-AT cells treated with the indicated doses of IMP-1088 or vehicle control (DMSO) at 1.5 hbi. (H, I) The quantification of the relative infection (H; number of N-positive cells normalized to number of nuclei) and mean intensity (I; a.u., arbitrary units) in A549-AT cells treated with indicated concentration of IMP-1088 or DMSO vehicle control 1.5 hbi and infected SARS-CoV-2 wuh for 8 h. (J, K) The quantification of the relative infection (J; number of dsRNA-positive cells normalized to number of nuclei) and mean intensity (K; a.u., arbitrary units) in A549-AT cells treated with indicated concentration of IMP-1088, DMSO vehicle control or 4OH-dynasore (30 µM) 1.5 hbi and infected SARS-CoV-2 wuh for 8 h. (L) Representative immunofluorescence images of human primary nasal epithelial cells grown as a pseudostratified columnar epithelium and treated with IMP-1088 (1 µM) or Nafamostat and Apilimod (20 µM and 0.25 µM, respectively) or DMSO vehicle control and infected with SARS-CoV-2 wuh for 72 h and stained against viral markers N (yellow), actin (magenta) and nuclei (cyan). (M) Representative image of haematoxylin and eosin-stained transfer section of an adult human nasal epithelium culture show scattered goblet cells (asterisks) and ciliated epithelial cells (arrowheads) of pseudo-stratified muco-ciliary epithelium. (N) The quantification of the relative infection (the number of N-positive cells normalized to the number of nuclei). (O) Representative immunofluorescence images of A549 cells treated with indicated concentrations of IMP-1088 or DMSO vehicle control at 2 hbi or 1 hpi of RSV-GFP (green, top panel), SFV-GFP (green, middle panel), and VSV-GFP (green, bottom panel) infection (P-R) Quantification of relative infection (number of GFP positive cells normalized to the number of nuclei) of RSV-GFP 24 hpi and 48 hpi (P), SFV-GFP at 12 hpi and 24 hpi (Q), and VSV-GFP at 12 hpi and 24 hpi (R), respectively, in DMSO and 0.01-1 µM IMP-1088 treated A549 cells (at 2 hbi or 1 hbi). Violin plots are median ± quartiles, line graphs are mean ± SEM. All results are from 3 biological replicates in each condition. Statistical testing was performed with one-way ANOVA with Tukey’s correction for multiple comparisons (E, H-J, N), two-way ANOVA with Dunnett’s multiple comparison test (B, C, F, P, Q, R) and unpaired t test (K).***p<0.001, ****p<0.0001, ns= non-significant.

At 48 hpi, the relative antiviral effect of IMP-1088 increased, with a reduction in infection by 80-90% compared to the DMSO control (Figs. 1D, E). In addition, the treatment prevented virus-induced cell-to-cell fusion (known as syncytia formation) (Fig. 1D: the enlarged inset i shows a multi-nucleated syncytium of cells, and ii/ iii show infected single cells). Next, to investigate the reversibility and minimum duration of IMP-1088 treatment sufficient to inhibit infections in cell cultures, A549-AT cells were pre-treated with either DMSO or 1 µM IMP-1088 for 1 to 120 minutes and, after the drug was removed by three washes, cell were infected with SARS-CoV-2 Wuh for 48 or 72 hours. The results revealed that IMP-1088’s inhibitory effect is virtually irreversible, and a one-minute treatment was sufficient to achieve a level of infection inhibition comparable to when the drug is continuously present in the media (Fig. S1A, B). This aligns with the extremely low dissociation constant reported for this drug (Kd < 210 pM)^19^. Notably, these results also demonstrated that at 72 hpi, SARS-CoV-2 Wuh relative infection was almost completely inhibited (∼98%) compared to the DMSO control (Fig. S1A, B). We also confirmed the antiviral effects of IMP-1088 against the SARS-CoV-2 variants Omicron and Delta, which showed similar inhibitory profile at 24 and 48 hpi as Wuh (Figs. 1F, G, and S1C, D).

To test whether the drug affected one of the ‘early’ stages of the viral life cycle, which include cell binding entry, viral protein synthesis, and genome replication, cells were pretreated with 0.1 µM or 1 µM IMP-1088 at 1.5 hbi, and infected with SARS-CoV-2 Wuh for 8 h. Quantification of relative infection at this early time point revealed that the drug-treatment had no noticeable impact on viral infection, indicating that the drug does not perturb viral entry (Figs. 1H and S1E), viral protein synthesis (Fig. 1I) or RNA replication (Fig. 1J). In contrast, a pre-treatment with an inhibitor of endocytosis, 4OH-dynasore, previously shown to prevent SARS-CoV-2 cell entry^19^, inhibited virus infection significantly at this time point (Figs. 1K, S1F). We next assessed the potential cytotoxic effects of IMP-1088 on A549-AT cells after 24 and 48 h of treatment by measuring cellular viability using CellTiter-Glo assay. Confirming previous findings^19^, we observed no significant effects on the cell viability compared to the vehicle control (Figs. S1G, S1H).

To assess whether the IMP-1088-induced replication defects lead to accumulation of viral genome mutations, we next compared the sequences of the SARS-CoV-2 Wuh isolates from DMSO and IMP-1088 treated cells to the first isolates of SARS-CoV-2 Wuh (NCBI Reference Sequence: NC_045512.2). At consensus level, the original SARS-CoV-2 Wuh sequence was identical to those isolated from the cells treated with DMSO or IMP-1088 (Table 1). Further, all minority variants in experimental samples were detected with low frequency (less than 5% of the virus population) and none of them seem to be specific for drug treatment.

Together, these findings indicate that the critical steps in the virus life cycle from entry to genome replication were not significantly affected by the drug treatment, suggesting that the antiviral effects of IMP-1088 were due to an interreference in a step of SARS-CoV-2 infection that follows genome replication.

### IMP-1088-treatment inhibits SARS-CoV-2 infection in human pseudostratified mucociliary nasal epithelial cells

SARS-CoV-2 enters the human body mainly through upper respiratory tract^25^. To investigate the antiviral efficacy of IMP-1088 in nasal cells, nasal epithelial cells from adult donors, which endogenously express ACE2 and TMPRSS2, were differentiated at the air–liquid interface as previously described^26^ until ciliated cells and mucus were observed (Fig. 1M), and cells obtained a transepithelial electrical resistance (TEER) measurement greater than 800 Ω/cm2^25^. Fully differentiated cultures were then treated with DMSO (vehicle control), 1 µM IMP-1088, or, as a positive antiviral control a combination of 20 µM Nafamostat and 0.25 µM Apilimod^27^ 1.5 hbi, infected with SARS-CoV-2 alpha at 10^6^ foci-forming units (FFU) per insert for 48 h, and then fixed and immunostained against N protein (Fig. 1L). Consistent with our findings in A549-AT cells, NMT inhibition strongly prevented the spreading of SARS-CoV-2 infection in human primary nasal cultures, while blocking virus entry by combined inhibition of TMPRSS2 using Nafamostat^28,29^ and endosome maturation using Apilimod^30,31^, resulted in a complete inhibition of infection (Fig. 1N). These results collectively demonstrate the effectiveness of IMP-1088 as an inhibitor of SARS CoV-2 spreading not only in cell lines but also in primary human nasal epithelial cells.

### IMP-1088 inhibits Respiratory Syncytial Virus but not Semliki Forest or Vesicular Stomatitis Virus infection

To date, the antiviral activity of IMP-1088 has been demonstrated against the common cold rhinovirus, vaccinia virus, and yellow fever virus which all harbour one or more proteins that undergo N-myristoylation^19–21^. Building on our results demonstrating that inhibition of host N-myristoylation using IMP-1088 inhibits SARS-CoV-2 infection, we expanded these experiments to other viruses that do not, based on current knowledge, express N-myristoylated proteins: Semliki Forest virus (SFV, family *Togaviridae*)^32^, Vesicular stomatitis virus (VSV, family *Rhabdoviridae*)^33^, and Respiratory syncytial virus (RSV, family *Pneumoviridae*)^34^. These three viruses were previously engineered to express green fluorescent protein (GFP) as a reporter gene. i.e. SFV-GFP, VSV-GFP, RSV-GFP^35–37^. A549 cells were pre-treated with DMSO vehicle control or with 0. 1 or 1 µM of IMP-1088 at 2 hbi or 1 hpi and infected with SFV-GFP, VSV-GFP, RSV-GFP at low doses (Fig. 1O). Image analysis revealed that IMP-1088 treatment inhibited the infection of RSV-GFP (MOI 0.2) by more than 50% compared to DMSO control at 48 hpi (Fig. 1P). SFV-GFP and VSV-GFP infections, on the other hand, were not affected by the drug treatment at 12 hpi (MOI 0.5), or 24 hpi (MOI 0.01) a time sufficiently long for these viruses to undergo multiple rounds of infection (Fig. 1Q, R). Together, our results demonstrate that pharmacological inhibition of host NMTs by IMP-1088 does not affect the early steps of the life cycle for any of the four viruses tested, and further indicates that the drug selectively impedes the infection of SARS-CoV-2 and RSV-GFP at later stages, namely assembly and/or release.

### IMP-1088 inhibits SARS-CoV-2 infectivity in human choroid plexus-cortical brain organoids, providing significant neuronal protection against the virus

SARS-CoV-2 has been shown to infect brain choroid plexus and to disrupt the blood-cerebrospinal fluid barrier^38,39^. To investigate the antiviral potential of IMP-1088 on SARS-CoV-2 Wuh infecting human choroid plexus, we employed a recently developed human choroid plexus-cortical organoid (ChPCOs) model^39^. This model is particularly advantageous for investigating choroid plexus and neurotropic viral infections, as previously demonstrated in our study that linked SARS-CoV-2 infectivity to neuronal cell death within cortical structures facilitated by the choroid plexus^40^. We generated ChPCOs and characterized their cellular composition at two months post-induction using single-cell RNA sequencing (scRNA-seq), which confirmed the presence of diverse neural cell types including cortical hem, choroid plexus, neural stem cells, radial glia, astrocytes, and neurons (Figs. 2A, S2-S4). Analysis of gene expression pertinent to SARS-CoV-2 entry and replication identified key receptors predominantly in choroid plexus and neuronal cells (Figs. 2B, 2C, S4). Two-month-old ChPCOs were then treated with DMSO, 1 µM IMP-1088, 0.25 µM Apilimod or 20 µM Nafamostat at 1.5 hbi and infected with SARS-CoV-2 Wuh for 72 h, and then fixed and immunostained for N protein, neuronal marker MAP2 and choroid plexus marker TTR (Fig. 2D). Compared to control organoids treated only with vehicle (DMSO), those ChPCOs treated with IMP-1088 or Apilimod exhibited significantly reduction in fewer SARS-CoV-2 infected cells (Figs. 2D, 2F). ChPCOs expressed no detectable levels of TMPRSS2 (Fig. 2B) and, therefore, the treatment with Nafamostat showed no effective inhibition of the SARS-CoV-2 infection in this model (Fig. 2F). Immunostaining further demonstrated a high susceptibility of neurons (MAP2 positive) to SARS-CoV-2 infection in two-month-old ChPCOs (Figs. 2D, 2G). Treatments with IMP-1088 and Apilimod markedly inhibited viral infection levels, while Nafamostat showed no effective inhibition of the SARS-CoV-2 infection (Figs. 2D, 2G). Both IMP-1088 and Apilimod treatments also had neuro-protective effects, as these treatments significantly decreased the fluorescence signal of immunodetected cleaved Caspase-3, a marker of apoptosis, in infected ChPCOs (Fig. 2H). In addition, Wwe also quantified the level of SARS-CoV-2 infection in ChPCOs using virions released from either DMSO or IMP-1088-treated cells (Fig. 2E). We observed significantly decreased levels of infection caused by virions released from IMP-1088-treated cells compared to those released from DMSO-treated cells (Figs. 2I-J). Collectively, these data further demonstrate that inhibition of cellular NMT1 via IMP-1088 inhibits SARS-CoV-2 infectivity and reduces the infectivity rate of its released virions.

**Figure 2.**
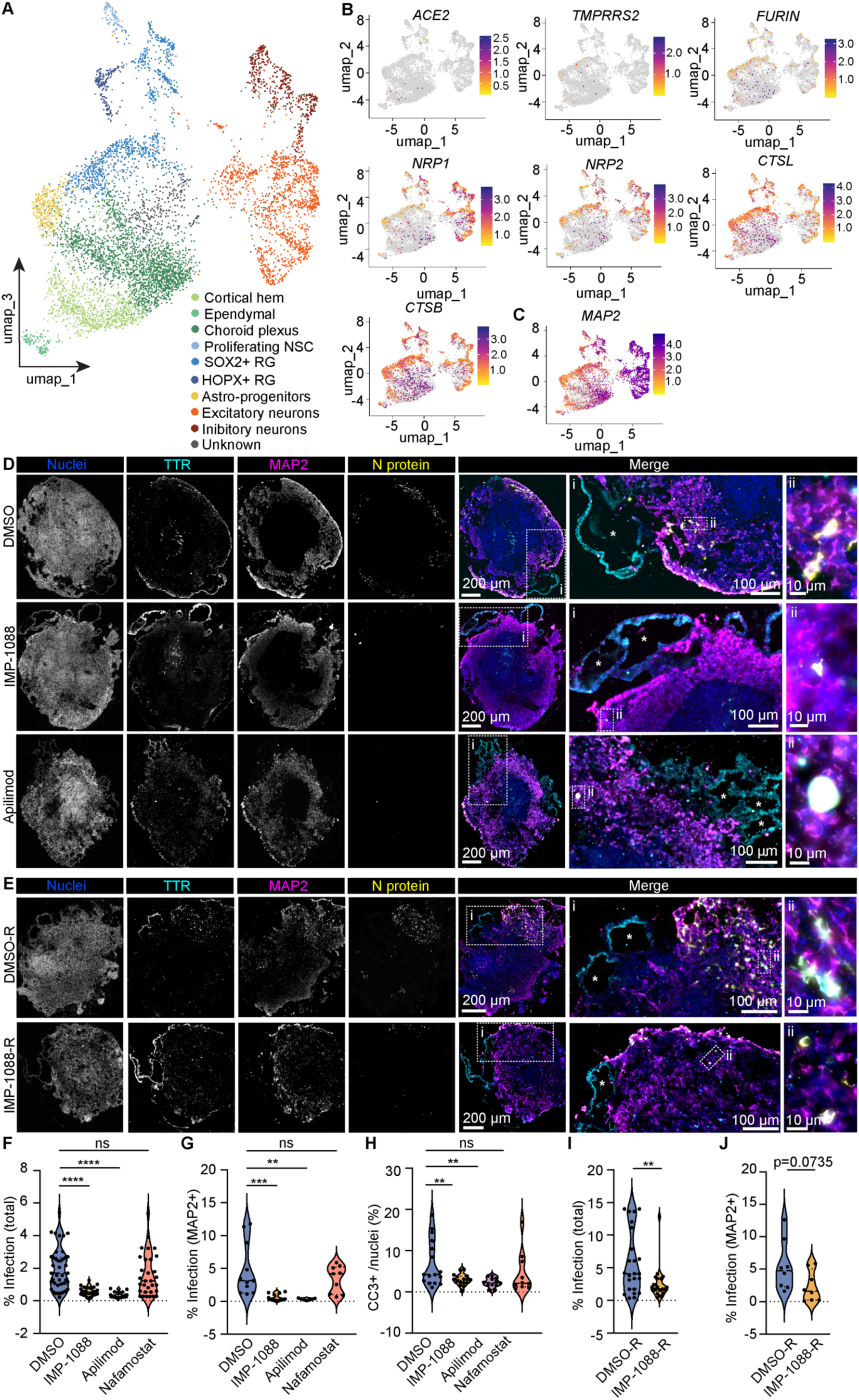
IMP-1088 inhibits SARS-CoV-2 infection in human choroid plexus-cortical brain organoids (ChPCOs). (A) Uniform Manifold Approximation and Projection (UMAP) visualisation of scRNA-seq on ChPCOs with cells coloured by cell types. (B, C) SARS-CoV-2 receptor and protease marker expression profile (B), mainly enriched in neuronal cells shown by MAP2 expression (C). Colour scale represents log transformed counts across cells (purple highly expressed, yellow less expressed). (D) Representative immunofluorescence images of ChPCOs treated with IMP-1088 (1 µM), Apilimod (0.25 µM) or DMSO vehicle control, 1.5 hours before-infection (hbi), infected with SARS-CoV-2 wuh for 72 h and stained against viral marker N (white), TTR (cyan), MAP2 (Magenta), and nuclei (blue). Boxed areas i and ii are shown magnified on right. (E) Representative immunofluorescence images of ChPCOs infected with 1×10^6 copy number of SARS-CoV-2 wuh released from IMP-1088 (IMP-1088-R) or DMSO (DMSO-R) treated A549-AT cells. Immunostaining shown as in D. (F, G) Quantification of the percentage of total infection (the number of N-positive cells normalized to the number of nuclei) in all cells, and in MAP-2 positive neuronal cells (G) in ChPCOs treated with DMSO (vehicle control), IMP-1088 (1 µM), Apilimod (0.25 µM), or Nafamostat (20 µM). (H) Quantification of the percentage of cleaved caspase 3 (CC3) positive cells normalized to the number of nuclei in ChPCOs treated with DMSO (vehicle control), IMP-1088 (1 µM), Apilimod (0.25 µM) or Nafamostat (20 µM). (I, J) Quantification of the percentage of total infection (H) and neuronal infection (MAP-positive, I) in ChPCOs using 1×10^6 copy number of SARS-CoV-2 wuh released from IMP-1088 (IMP-1088-R) or DMSO (DMSO-R) treated A549-AT cells (as shown in D). Violin plots are median ± quartiles. Results are from 2 technical replicas, dots present immunostained sections. Statistical testing was performed with one-way ANOVA with Kruskal-Wallis multiple comparison test (F), ordinary one-way ANOVA multiple comparison test (G), two-way ANOVA multiple comparison test (H), Mann-Whitney test (I) and unpaired t test (J). **p<0.01, ***p<0.001, ****p<0.0001, ns= non-significant.

### Inhibition of host NMTs does not interfere with the release, but impairs the infectivity, of SARS-CoV-2 progeny virions

Our results from ChPCOs infected with virions released from IMP-1088-treated cells (Fig. 2E, 2H, 2I) suggest that the progeny virions released from NMT inhibited cells are less infectious than those released from control cells. To gain deeper insights into this, we next investigated whether the IMP-1088-treatment affected viral genome release using quantitative real-time PCR (qRT-PCR) and primers targeting a specific region of the viral RNA that encodes the RNA-dependent RNA polymerase (RdRp). In short, A549-AT cells were pretreated with either DMSO as a control or IMP-1088, then infected with SARS-CoV-2 Wuh for 2 hours. Afterward, the cells were washed three times with PBS to remove any residual virus from the media and replaced with fresh media, with or without the drug. Culture media was collected at 0 h (immediately after media replacement), 6 h, 12 h, 24 h, and 48 h post-media replacement and analysed by qRT-PCR. The viral RNA copy number in the media collected from cells treated with media with or without IMP-1088 showed no significant differences up until 12 hours after media replacement (Figs. 3A, 3B). However, 24 hours after media replacement, and even more so at 48 hours, the number of virions (measured by the amount of viral RNA present in the extracellular media) released from IMP-1088-treated cells dropped significantly, decreasing 10-fold by 48 hours compared to virions released from DMSO-treated cells (Figs. 3A, 3B). Given that no significant differences were observed up until 12 hours post-media replacement, and that the 24 h and 48 h time points allow multiple rounds of cell-to-cell infection, these results indicate that IMP-1088 treatment does not have inhibitor effect on viral release. However, the lower levels of extracellular viral RNA detected at 48 hours after media replacement in IMP-1088-treated cells could indicate a reduction in the released virion infectivity impairing cell-to-cell transmission.

**Figure 3.**
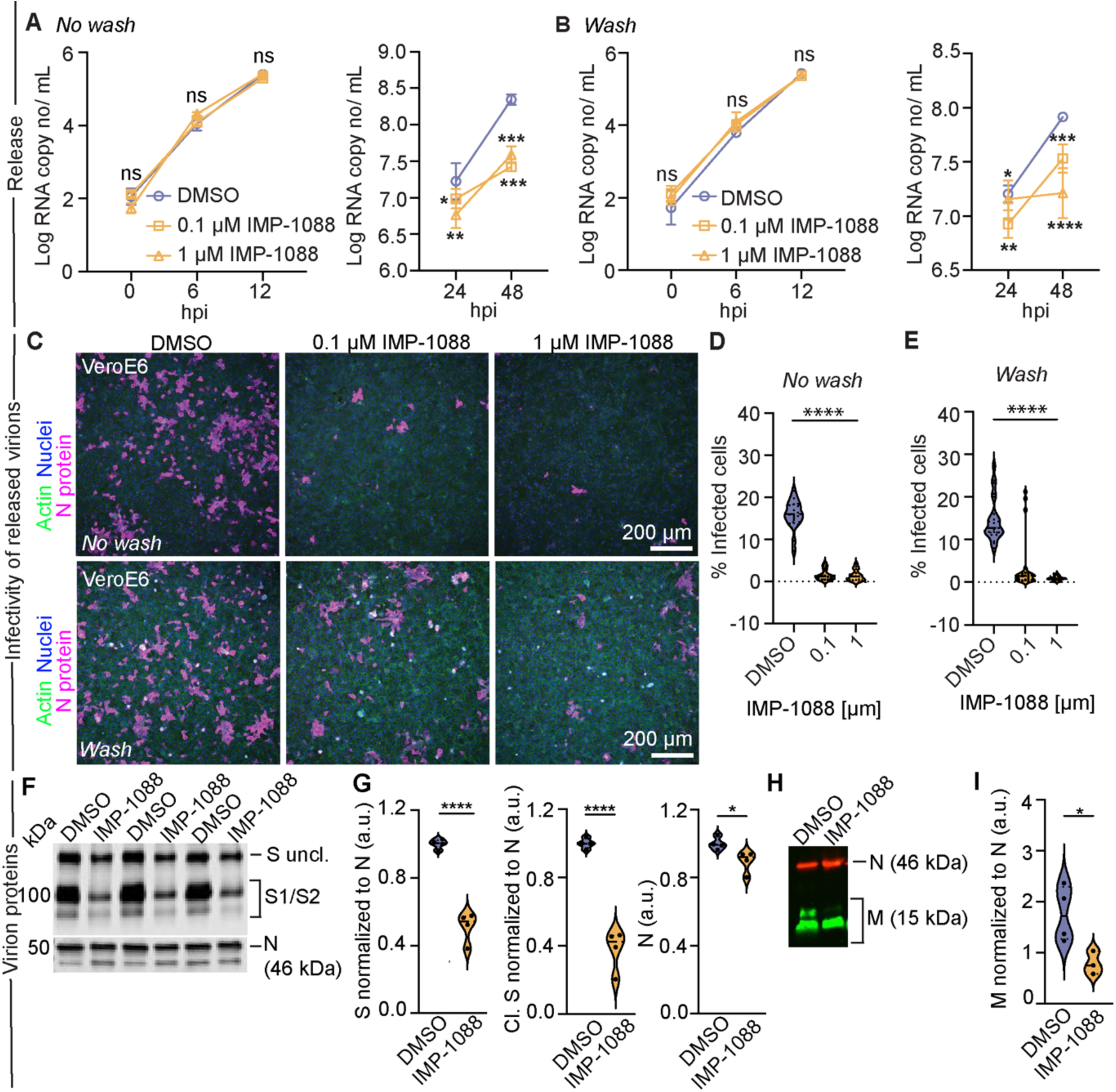
IMP-1088 treatment does not impair the release of SARS-CoV-2 but significantly compromises the re-infectivity of the released virions by impairing spike incorporation and maturation. (A, B) The qRT-PCR quantifications of the released SARS-CoV-2 wuh from A549-AT cells treated with indicated concentrations of IMP-1088 or DMSO vehicle control at 2 hbi and infected with SARS-CoV-2. At 2 hpi, the infection media was washed three time with PBS and replaced with new media with indicated concentrations of IMP-1088 (no wash, A) or without IMP-1088 (wash, B). The media was collected at 0, 6, 12, 24, and 48 hpi from infected cells and the SARS-CoV-2 wuh RNA copy numbers were quantified using qRT-PCR. (C) Representative immunofluorescence images of VeroE6 cells infected with inocula containing the same number of viral RNA genomes of SARS-CoV-2 wuh and immunostained for N protein (magenta), actin (green), and nuclei (blue). The viruses were isolated from A549-AT cells pre-treated with indicated IMP-1088 concentrations or DMSO vehicle control at 2 hbi. Drugs were either washed out (top panel) or kept on cells for the duration of the experiment (bottom panel). (D, E) Quantification of the relative infection of SARS-CoV-2 wuh of released from A549-AT cells treated with 0.1 µM or 1 µM IMP-1088 for the duration of the experiment (D) or following a wash out (E) at 24 hpi in VeroE6 cells. (F) Western blot of uncleaved viral spike (S) ± glycosylation and N protein of the same copy number of purified SARS-CoV-2 virions released from A549-AT cells treated with DMSO or 1 µM of IMP-1088 for 48 h. (G) Western blot quantification of S protein and cleaved (cl.) S protein levels normalized to N protein, and N protein levels. (H) Western blot of viral M and N proteins of the same copy number of purified SARS CoV-2 virions released from A549-AT cells treated with DMSO control or 1 µM of IMP-1088 for 48 h. (I) Quantification of viral M protein expression normalized to N protein from western blots. Violin plots are median ± quartiles. Results are from 3 biological replicates in each condition. Statistical testing was performed with ordinary one-way ANOVA multiple comparison test (A, B, D, E), and unpaired t test (G-I). *p<0.05, **p<0.01, ***p<0.001, ****p<0.0001, ns= non-significant.

**Figure 4.**
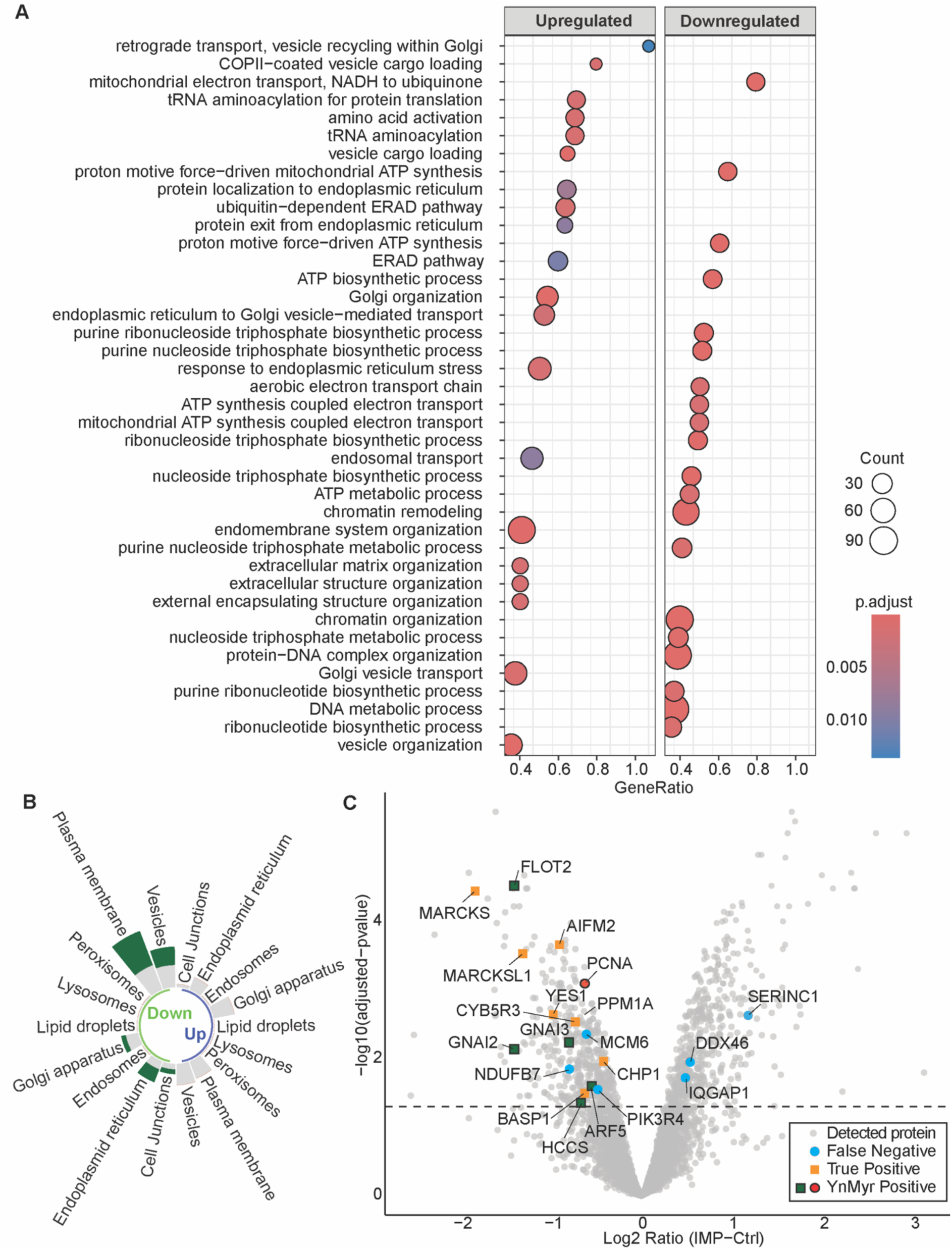
Quantitative mass spectrometry analysis reveals NMT inhibition alters the host proteome. (A) Gene Ontology (GO) pathway enrichment analysis of A549-AT cells treated with 1 µM of IMP-1088 or DMSO control for 48 h and analysed by label-free mass spectrometry show subcellular components on Y axis and gene ratio of upregulated and down regulated protein on X axis with IMP-1088 treatment (DA adjust p≤0.05). (B) GO analysis of ‘endomembrane system’ term – using protein atlas subcellular location. Split into Up and Down (abundance) and stacked with NMT positive targets (green). (C) Volcano Plot shows all detected proteins from IMP-1088-treated cells compared to DMSO-treated control cells and to experimental data from HeLa cells treated with IMP-1088 (Kallemeijin et al., 2019). False negative (blue; experimentally supported by Kallemeijin et al., 2019 but not containing an NMT motif), true positive (orange; predicted by NMT motif and experimentally supported by Kallemeijin et al., 2019) and YnMyr positive targets (NMT targets, green rectangles and red circles with black outlines). Predicted refers to inferred by NMT containing motif only. Odds Ratio of decreased versus increased abundance: 10.36982, Fisher Test p-value 6.009e-05) confirming previously reported *N*-myristoylated proteins (grey outlines, Mousnier *et al.* 2018). N=5 biological replicates in each condition.

To explore this further, we investigated whether virions released from IMP-1088-treated A549-AT cells were less infectious compared to those released from cells treated with DMSO. A549-AT cells were pretreated with either DMSO or IMP-1088, then infected with SARS-CoV-2 Wuh strain. At 2 hpi, the cells were washed three times with PBS to remove any residual virus from the media, then cultured in fresh media with or without the drug for 48 h. Culture media containing virions was collected at 48 hours post-infection, and the viral RNA copy numbers in each sample were quantified by qRT-PCR. VeroE6 cells were then inoculated with media containing equal amounts of SARS-CoV-2 Wuh RNA, representing virions released from either IMP-1088-treated or DMSO-treated (± drug wash) A549-AT cells, and at 24 hpi cells were fixed and immunostained against N protein (Fig. 3C). Quantification of the percentage of infection revealed that virions released from IMP-1088-treated cells were significantly less infectious (∼90%) compared to those from DMSO-treated controls (Figs. 3D, 3E). This reduced infectivity was observed across all IMP-1088 treatments, regardless of whether the drug was washed off or remained in the culture media for the duration of the experiments. Taken together, these findings suggest that inhibiting host NMT with the small molecule inhibitor IMP-1088 does not significantly impact the early stages of SARS-CoV-2 infection or virion release, but rather it diminishes the infectivity of newly released virions.

### Host NMT activity is crucial for SARS-CoV-2 progeny virion S protein composition and cleavage

Following our findings of host cell NMT inhibition leading to a significant decrease in infectivity of the released virions and the perturbed trafficking of the viral particles to ERGIC and Golgi compartments, we next explored the potential impact of this transport blockage on the structural proteins of the released virions. We isolated SARS-CoV-2 virions from A549-AT cells treated with DMSO or IMP-1088 and conducted western blot analyses, normalizing by the same viral RNA copy number. We observed slightly decreased levels of the viral N protein, which is important for viral genome packaging, in virions released from both DMSO and IMP-1088-treated cells (Figs. 3F, 3G). In contrast, analysis of the S protein, which is key to the viral infectivity, indicated that virions released from IMP-1088-treated cells contained significantly less total S protein, which also showed significantly reduced levels of glycosylation and cleavage compared to virions released form DMSO-treated cells (Figs. 3F, 3G). The reduced spike expression aligned with a decreased expression of M protein, essential for viral assembly in the ERGIC, which also exhibited lower levels and reduced glycosylation compared to DMSO control (Figs. 3H, 3I). These results indicate that the loss of SARS-CoV-2 progeny virions infectivity following treatment with IMP-1088, results from perturbed assembly of the virions with decreased levels of S protein.

### NMTs regulate protein homeostasis

To understand why the released virions were less infectious, we characterized the proteomic changes induced by NMT inhibition. We conducted global proteome analysis using label-free mass spectrometry on A549-AT cells treated with DMSO or IMP-1088 for 48 h. Replicate reproducibility of the analysis was assessed by principal component (Fig. S5A) and hierarchical clustering (Fig. S5B) analysis, confirming that the proteome identified in each replicate consistently clustered according to its respective experimental group (i.e., A549-AT +DMSO and A549-AT +IMP-1088). The inhibition of cellular NMT led to significant changes in the abundance of proteins across multiple subcellular pathways (Fig. 4A), particularly those involved in the early secretory pathway endoplasmic reticulum (ER), ER-Golgi intermediate compartment (ERGIC), and Golgi apparatus and vesicle trafficking (Figs. 4B). Further analysis revealed a significant decrease in the abundance of proteins with N-terminal glycine destined for *N*-myristoylation (Fig. 4C; Table 2^24^). Together, our findings suggest that IMP-1088 alters host cell protein, especially *N*-myristoylated protein, abundances and particularly those related to the early secretory pathway and intracellular membrane trafficking. These finding support previous literature suggesting that host cell NMT inhibition leads to degradation of *N*-myristoylated proteins, as myristylation is crucial for e.g. protein stability and function^41,42^.

### NMTs control the structure and dynamics of the early secretory pathway

Following the findings of the mass spectrometry analysis, we next investigated how changes in protein homeostasis might affect the ultrastructure of the ER, ERGIC, and Golgi using electron microscopy (EM). We observed distinct morphological changes in early secretory compartments following treatment with IMP-1088. In control cells, the ER appears as a network of interconnected membrane-enclosed compartments extending from the nuclear envelope (NE) into the cytoplasm (Fig. 5A). In IMP-1088-treated cells, NE and the ER network exhibited noticeable dilation at 24 hours of treatment (Fig. 5B), with further enlargement at 48 hours (Fig. 5C). Similar dilation of the organelle lumen was also observed in ERGIC and Golgi apparatus in IMP-1088-treated cells (Figs. 5A-C), indicating a perturbation of membrane trafficking in the secretory pathway.

**Figure 5.**
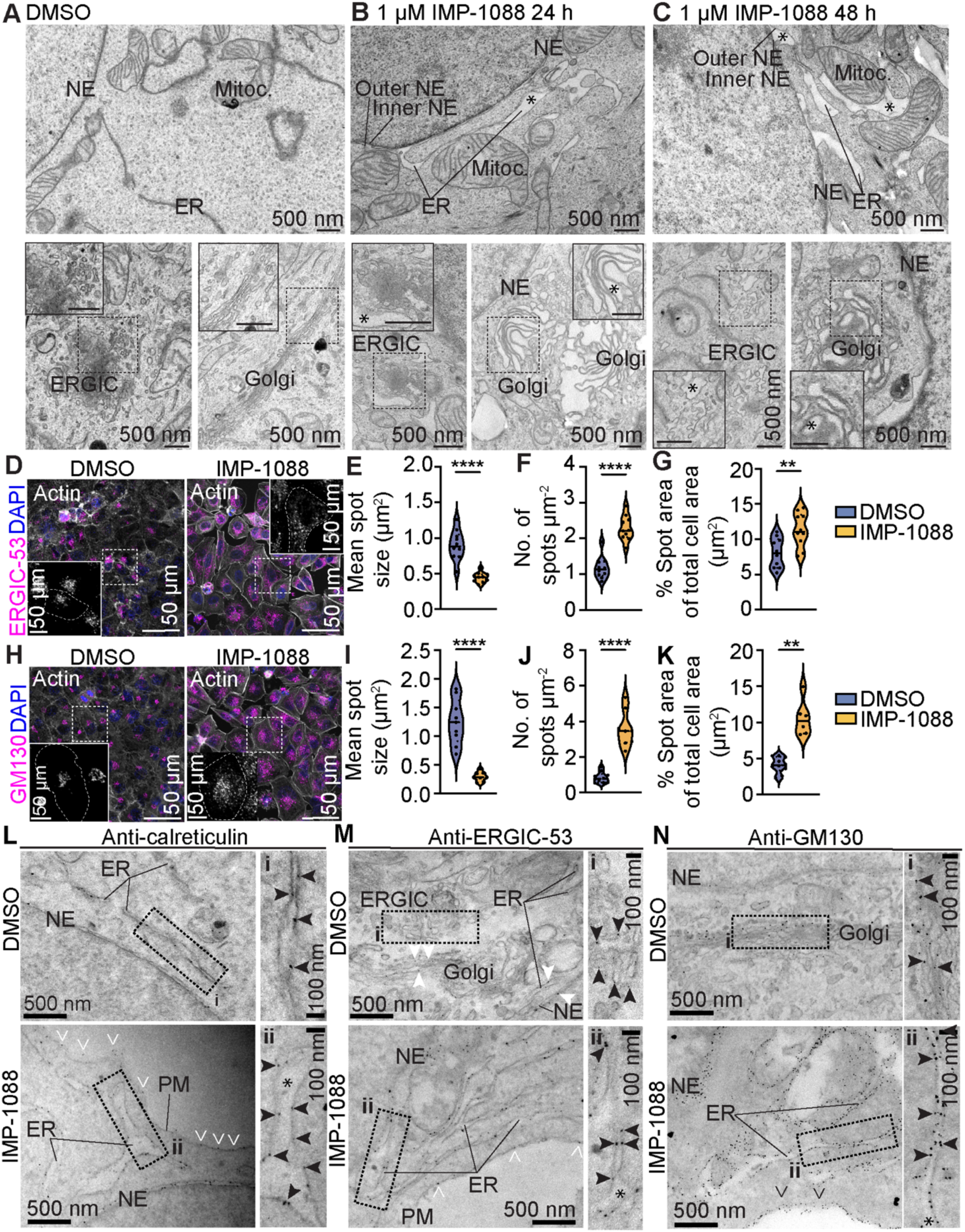
IMP-1088 alters ERGIC and Golgi apparatus ultrastructure. (A, B, C) Representative electron microscopy (EM) images of A549-AT cells treated with DMSO vehicle control (A), 1 µM of IMP-1088 for 24 h (B) or 48 h (C) show ultrastructure of rough endoplasmic reticulum (RER), ERGIC, Golgi, nuclear envelope (NE), mitochondria (Mitoc.) and dilated luminal space (asterisks). Boxed areas are shown in corners with higher magnification. Bars 500 nm. (D) Representative immunofluorescence images of A549-AT cells treated with 1 µM of IMP-1088 or DMSO vehicle control for 48 h and stained against endogenous ERGIC-53 (magenta) and nuclei (blue). Boxed areas are shown on right with higher magnification. (E-G) Quantifications of ERGIC-53 average spot size (µm^2^) (E), number of spots per µm^2^ (F), and percentage of spot area of the total cell area (G) in indicated conditions. (H) Representative immunofluorescence images of A549-AT cells treated with 1 µM of IMP-1088 or DMSO vehicle control for 48 h and stained against endogenous GM130 (magenta) and nuclei (blue). Boxed areas are shown on right with higher magnification. (I-K) Quantifications of GM130 average spot size (µm^2^) (I), no of spots per µm^2^ (J), and percentage of spot area of the total cell area (K) in indicated conditions. (L-N) Representative EM images of immuno-gold labelling (arrowheads) against endogenous calreticulin (L), ERGIC-53 (M) and GM130 (N) in A549-AT in indicated conditions. Plasma membrane (PM) staining (white open arrowheads indicate extracellular staining, while black open arrowheads indicate PM staining on cytosolic side), endoplasmic reticulum (ER), nuclear envelope (NE) and dilated luminal space (asterisks) are indicated. White arrowheads in (M) indicate ERGIC-53 staining at Golgi, ER and NE. Boxed areas i-ii are shown with higher magnification on right. Violin plots are median ± quartiles. A-G results are from 3 biological replicates in each condition, H-J from one. Statistical testing was performed with Mann-Whitney test (E) and unpaired t test (F, G, I-K). **p<0.01, ****p<0.0001.

Consistent with our EM data, immunostaining against endogenous ERGIC-53, which is a mannose-specific membrane lectin operating as a cargo receptor for the transport of glycoproteins from the ER to the ERGIC^43^, showed a compact organelle structure in the perinuclear region in DMSO-treated A549-AT cells (Fig. 5D). In contrast, treatment with IMP-1088 caused dispersal and fragmentation of the ERGIC-53 signal (Fig. 5D). Quantification of the ERGIC-53 immunofluorescent staining revealed a significant decrease in average spot size, increase in number of spots and area covered by the spots (Fig. 5E). Similar results were obtained from immunofluorescent staining against endogenous protein GM130^44^, which is a peripheral cytoplasmic protein that is tightly bound to Golgi apparatus (Fig. 5F, 5G).

To characterize these phenotypes further, we performed immuno-EM analysis of endogenous calreticulin, ERGIC-53 and GM130 in A549-AT cells treated with DMSO or IMP-1088 for 48 h. Calreticulin staining was mainly observed in ER and NE lumen in DMSO treated cells (Fig. 5H). Following IMP-1088 treatment, the calreticulin staining was observed in the dilated ER and NE lumen (Fig. 5H). ERGIC-53 staining was mainly observed in ERGIC membranes, with some signal in Golgi and ER membranes, consistent with its function in membrane trafficking (Fig. 5I). IMP-1088 treatment caused a redistribution of ERGIC-53 signal to ER and NE membranes (Fig. 5I). GM130 signal was mainly observed on Golgi membranes in DMSO control cells (Fig. 5J). IMP-1088-treatment, on the other hand, caused a substantial relocation of GM130, which was found tightly tethered to ER membranes (Fig. 5J). It is noteworthy that we also observed calreticulin and ERGIC-53 signal outside of the cells, at the plasma membrane (Fig. 5H-5I). GM130 signal, on the other hand, was found tethered to the cytoplasmic side of the plasma membrane (Fig. 5J).

N-myristoylation is a conserved feature of ARFs, including ARF1^45^. To characterize the effects of IMP-1088 treatment on secretory pathway trafficking further, we next compared the of IMP-1088 treatment to Brefeldin-A (BFA), which is an ADP-ribosylation factor 1 (ARF1) inhibitor^46^ that blocks protein transport from ER to the Golgi complex by preventing association of COP-I coat to the Golgi membranes, and triggering a rapid distribution of Golgi proteins to ER^47^. For this, we performed live cell imaging of A549-AT cells transiently expressing ARF1-GFP^48^. In control cells, ARF1 signal was found at the perinuclear area localizing to Golgi apparatus (Fig. S6A). A treatment with 5 µg/mL BFA for 40 s or 4 min led to a loss of the Golgi localization and to a homogenous distribution throughout the cell (Fig. S6A,B). These effects were different from IMP-1088, where we discovered that the treatment with 1 µM IMP-1088 for 1h or 48 h induced a gradual dispersal and fragmentation of Arf1-GFP (Fig. S6C; Note the considerably longer treatment). We observed a gradual increase in the mean fluorescence intensity (Fig. S6D), along with decrease in the mean spot size (Fig. S6E), and an increase in the number of spots (Fig. S6F), of ARF1-GFP following IMP-1088 treatment. Together, these results indicate that the effects of IMP-1088 to the secretory pathway organelle morphology and trafficking are different from BFA, which is a well-established inhibitor of ARF1.

### Inhibition of NMTs perturbs SARS-CoV-2 intracellular trafficking

We next asked how does the observed perturbed morphology of ER, ERGIC and Golgi affect trafficking of SARS-CoV-2 in the secretory pathway. In short, A549-AT were treated with DMSO or IMP-1088 1.5 hbi, infected with SARS-CoV-2 Wuh for 48 hours, and then fixed and processed for EM. We observed abundant viral particles in the lumen of nuclear envelope and ER and in ERGIC and Golgi compartments (Fig. 6A). Viral particles were also observed inside of convoluted membranes (CMs) and large membrane-bound vesicles (LVCVs). In IMP-1088-treated cells, we observed a dilation of ER, ERGIC and Golgi apparatus lumens (Fig. 6B) consistent with our previous results (Figs. 5B, 5C). Although viral particles were observed in the ER, CMs and LCVSs, they were notably absent in ERGIC and Golgi structures (Fig. 6B), suggesting a transportation blockage of the virus from ER to ERGIC and Golgi. Additionally, we observed in control A549-AT cells that virions were released from large vacuoles, while in IMP-1088 cells we observed an additional virus-containing membrane clusters fusing with the plasma membrane and forming small tunnels for the viral egress (Figs. 6A, 6B). The origin of these membrane clusters is currently unknown.

**Figure 6.**
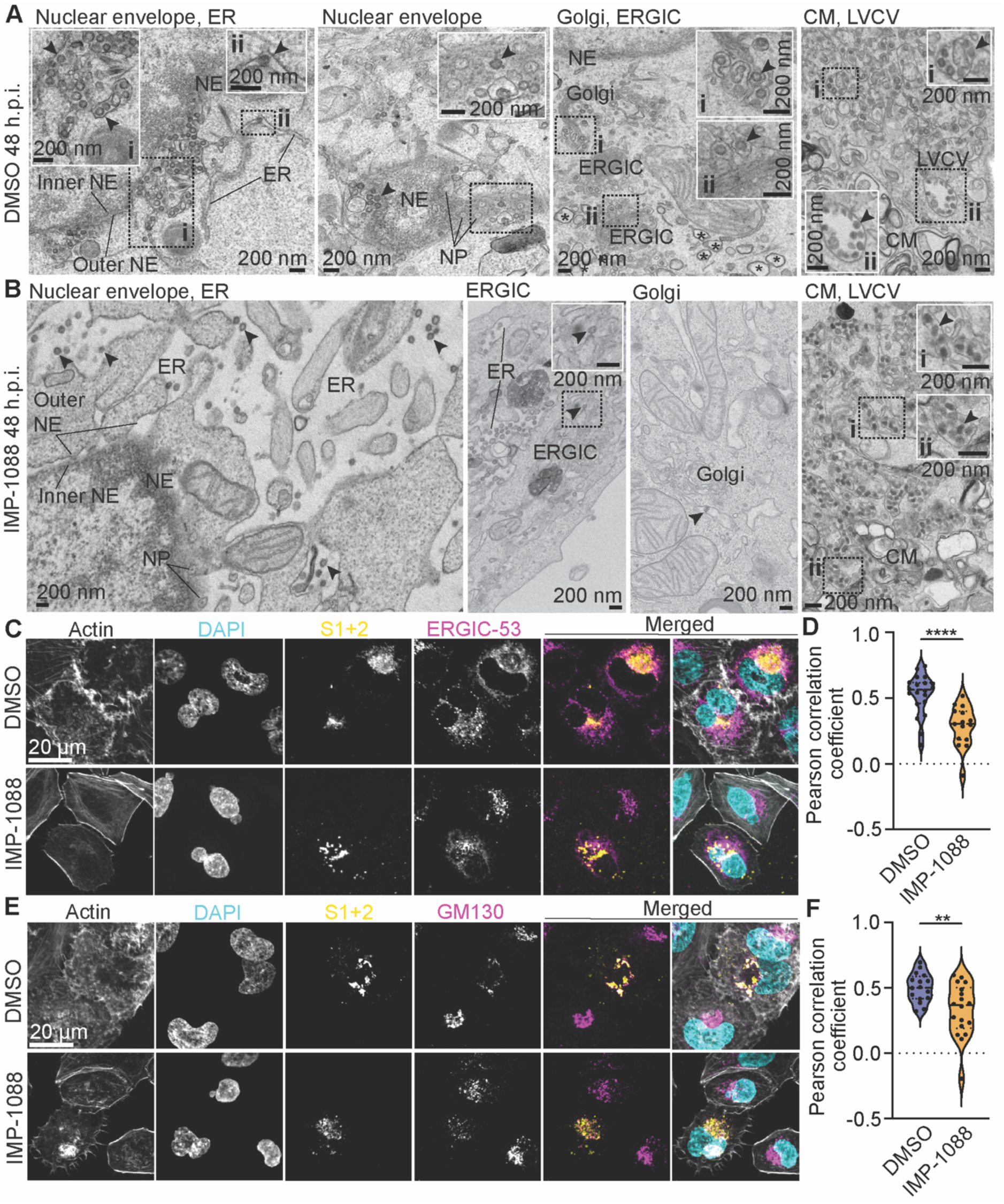
Inhibition of host NMTs leads to SARS CoV-2 egress via ERGIC-and Golgi-independent secretory pathway. (A, B) Representative EM images of A549-AT cells treated with DMSO vehicle control (A) or 1 µM of IMP-1088 (B) at 2 hbi and infected with SARS CoV-2 wuh for 48 h showing viral particles (arrowheads) in endoplasmic reticulum (ER), ERGIC, Golgi, nuclear envelope (NE), clustered membranes (CM), and large vesicle containing vesicle (LVCV). Boxed areas i and ii are shown with higher magnification in corners. (C) Representative immunofluorescence images of A549-AT cells treated with 1 µM of IMP-1088 or DMSO vehicle control 2 hbi, infected with SARS-CoV-2 wuh for 48 h and stained against endogenous ERGIC-53 (magenta), viral S (S1+2, yellow), actin (white) and nuclei (DAPI, cyan). (D) Quantifications of Pearson correlation coefficient of SARS-CoV-2 wuh S with ERGIC-53 in DMSO control and IMP-1088 treated cells. (E) Representative immunofluorescence images of A549-AT cells treated with 1 µM of IMP-1088 or DMSO vehicle control, infected with SARS CoV-2 wuh for 48 h and stained against endogenous GM130 (magenta), viral S (S1+2, yellow), actin (white) and nuclei (cyan). (F) Quantifications of Pearson correlation coefficient of SARS CoV-2 wuh S with GM130 in DMSO control and IMP-1088 treated cells. Violin plots are median ± quartiles. All the results are shown from 3 biological replicates in each condition. Statistical testing was performed with unpaired t test (D, F). **p<0.01, ****p<0.0001.

Supporting our EM findings, immunofluorescence analysis of SARS-CoV-2 S protein co-stained with endogenous ERGIC-53 (Fig. 6D, 6E) or GM130 (Fig. 6F, 6G) in A549-AT cells treated with DMSO or IMP-1088 and infected with SARS-CoV-2 for 48 hours, demonstrated that inhibition of NMTs decreased the co-localization of the viral S proteins with both ERGIC and Golgi apparatus, which are key cellular compartments needed for the proper viral assembly and maturation. We also observed consistent changes in the appearance of F-actin following IMP-1088 treatment (Fig. 6D, 6F; Note peripheral stress fibres and loss of shorter F-actin.). Together, these findings suggest that IMP-1088-treatment alters the host cell early secretory pathway structure and dynamics, perturbing viral assembly and maturation.

### Inhibition of host NMTs activates a Golgi-bypassing SARS-CoV-2 secretory pathway through ER and lysosome egress

Following our results collectively demonstrating that inhibition of host NMTs disrupted viral assembly and maturation in the Golgi, resulting in released virions with compromised infectivity, we next studied whether treating the virions released from IMP-1088-treated cells with Furin, which aids in spike protein cleavage in Golgi, could restore their infectivity. Furin treatment significantly increased the infectivity of virions released from IMP-1088-treated cells but had no significant effect on those from DMSO-treated controls (Fig. 7A).

**Figure 7.**
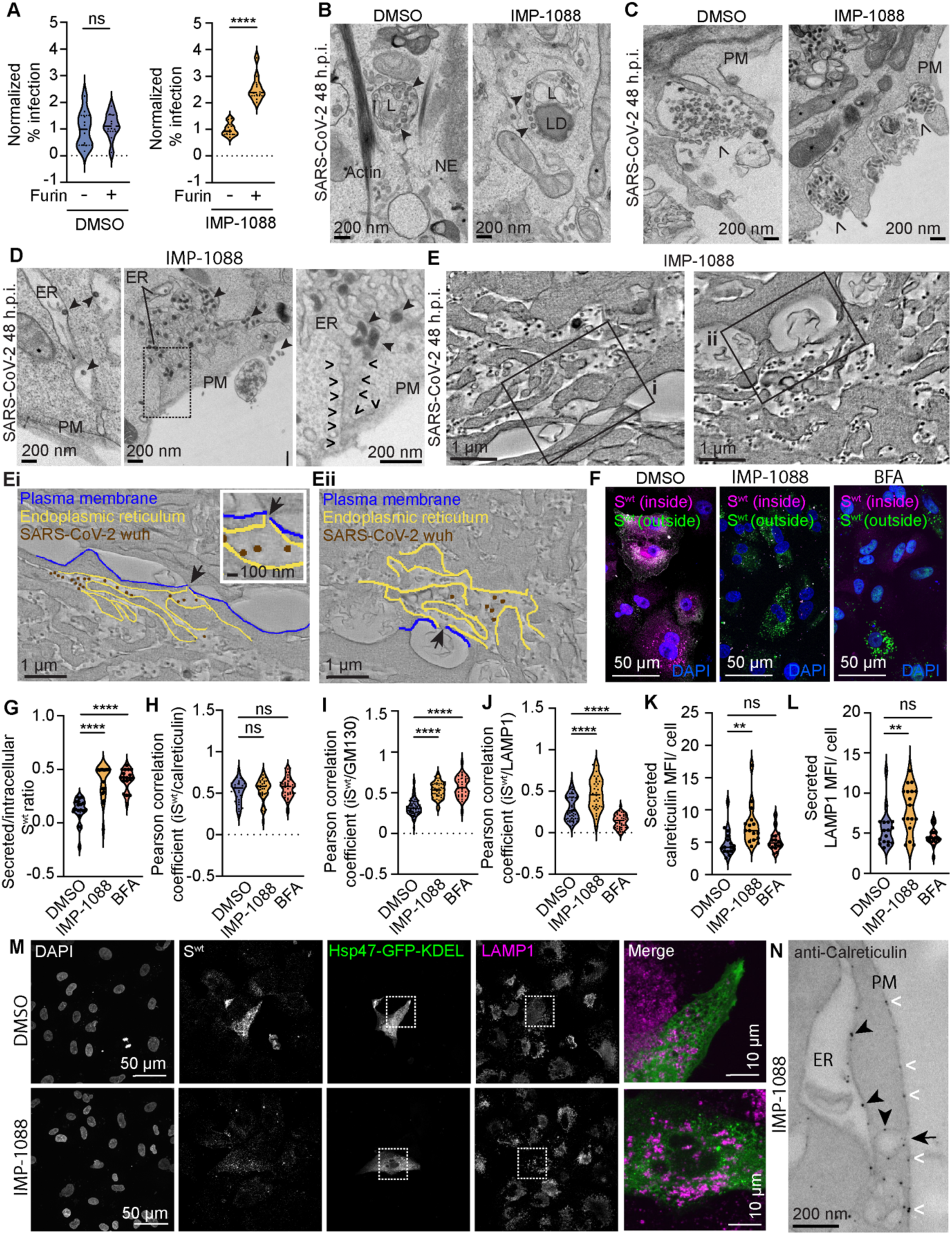
Inhibition of host NMTs leads to SARS-CoV-2 egress from lysosomal and ER compartments. (A) Quantification of the relative SARS-CoV-2 wuh infection (number of viral N positive cells normalized to the number of nuclei, DAPI) in VeroE6 cells infected with SARS-CoV-2 virions released from DMSO or 1 µM of IMP-1088 treated A549-AT cells, with and without a treatment with furin. (B) Representative EM images of A549-AT cells treated with DMSO control or 1 µM of IMP-1088 and infected with SARS-CoV-2 wuh (arrowheads) for 48 h showing viral particles in lysosomes (L). Actin bundle, lipid droplet (LD) and nuclear envelope (NE) are indicated. (C) Representative EM images of SARS-CoV-2 wuh egress (open arrowheads) from A549-AT cell plasma membrane (PM) in indicated conditions. (D) Representative EM images of SARS-CoV-2 wuh (arrowheads) in the lumen of endoplasmic reticulum (ER) adjacent to plasma membrane or opening to extracellular space (open arrowheads) in IMP-1088 treated A549-AT cells. The boxed area is magnified on right. (E) ET capture of an A549-AT cell treated with 1 µM IMP-1088 and infected with SARS-CoV-2 wuh for 48 h. Boxed areas i and ii are shown below with the respective segmentation and model showing ER (yellow) openings (arrows) at plasma membrane (blue) and viral particles inside ER (brown). Both regions of interest originate from the same tomogram, see Video 1. (F) Representative immunofluorescence image of A549-AT transiently expressing S^wt^, treated with DMSO, 1 µM IMP-1088 for 48 h, or 5 µg/µl brefeldin A (BFA) for 25 min and immunostained against internal (magenta) and secreted (green) S^wt^, followed by DAPI staining. (G) Mean fluorescence intensity (MFI) ratio of transiently expressed S^wt^ comparing secreted versus intracellular S^wt^ MFI in A549-AT cells in indicated conditions. (H-J) Pearson correlation coefficient of transiently expressed intracellular S^wt^ (iS^wt^) with calreticulin (H), GM130 (I) and LAMP1 (J), and in A549-AT cells treated with DMSO control, 1 µM of IMP-1088 for 48 h or 5 µg/µl BFA for 25 min. (K-L) MFI of secreted calreticulin (K) and LAMP1 (L) normalized to nuclei (DAPI) in A549-AT cells in indicated conditions. (M) Representative immunofluorescence images of A549-AT transiently expressing Hsp47-GFP-KDEL (green) and S^wt^, treated with DMSO control or 1 µM of IMP-1088 for 48 h, and immunostained against endogenous LAMP1 (magenta), and S protein, followed by DAPI staining. (N) Representative EM image of immuno-gold labelling (arrowheads) against endogenous calreticulin in IMP-1088-treated A549-AT cells. Plasma membrane (PM), endoplasmic reticulum (ER), ER opening to extracellular space (arrow), and extracellular staining (white open arrowheads) are indicated. Violin plots are median ± quartiles. All the results are shown from 3-6 biological replicates in each condition. Statistical testing was performed with unpaired t test (A) and ordinary one-way ANOVA multiple comparison test (G-L). **p<0.01, ****p<0.0001, ns= non-significant.

We next investigated the egress of the virions. SARS-CoV-2 has previously been shown to exploit lysosomal organelles for egress^49^. Based on our EM analysis, SARS-CoV-2 virions were observed in lysosomal structures both in DMSO controls and following IMP-1088 treatment (Fig. 7B). We next investigated whether these lysosomes presented endo-lysosomal compartments containing endocytosed SARS-CoV-2 or lysosomes involved in the egress of newly synthetised virions. In both DMSO control and IMP-1088-treated cells, SARS-CoV-2 virion egress was often observed to occur form large vacuole structure (Fig. 7C). Unexpectedly, in IMP-1088 treated cells, we also observed the release of virions from structures resembling ER (Fig. 7D). To investigate this further, we next studied the trafficking of SARS-CoV-2 S protein. For this, A549-AT cells were transfected with a plasmid expressing S^wt^, which, in the absence of other viral components, is retained in the secretory pathway (mainly ER and Golgi) and not secreted outside of the cells (Fig. 7E). Co-localization analysis of the transiently expressed S^wt^ against endogenous LAMP1 by immunofluorescence staining demonstrated that IMP-1088-treatment increased the localization of the S^wt^ with lysosomes, while 5 µg/µm BFA-treatment for 25 min, which blocks protein transport from ER to the Golgi complex by preventing association of COP-I coat with the Golgi membranes, and triggering a rapid distribution of Golgi proteins to ER^47^ (Fig. 7F). Co-localization analysis of transiently expressed S^wt^ immunofluorescence staining against endogenous calreticulin, luminal chaperone of ER, remained unaltered following both IMP-1088 and BFA treatments (Fig. 7G), while co-localization of S^wt^ with GM130 increased significantly in both treatments (Fig. 7H), likely reflecting the observed relocation of GM130 tethering from Golgi apparatus to ER membranes in IMP-1088 treated cells (Fig. 5N), and previously demonstrated disassembly of the Golgi complex and accumulation of secretory proteins in the ER following BFA-treatment^50^. Co-localization analysis of S^wt^ with endogenous LAMP1 demonstrated that IMP-1088-treatment increased the localization of the S^wt^ with lysosomes, while BFA-treatment decreased it (Fig. 7I). IMP-1088 treatment also led to increased secretion of endogenous calreticulin (Fig. 7J) and LAMP1 (Fig. 7K) detected outside cells. We observed no intermixing between luminal ER marker HSP47-GFP-KDEL^51^ and endogenous LAMP1 following IMP-1088 treatment (Fig. 7I), indicating the two organelles detain their structural identity^52^. Direct fusion of ER with plasma membrane in IMP-1088 treated A549-AT cells was detected using immuno-EM of endogenous calreticulin (Fig. 7M). Together these results indicate that the egress of virions in IMP-1088 treated cells occurs through ER and lysosomal carriers.

## Discussion

SARS-CoV-2 pandemic was declared over 2022 but COVID-19 continues to pose a global health issue. Current antiviral therapies against SARS-CoV-2 include systemically delivered mRNA vaccines^53–55^, viral vector vaccines^56,57^, small molecules that interfere with viral RNA synthesis (e.g., remdesivir, and molnupiravir)^58–62^, and viral protease inhibitors (e.g., Paxlovid)^63,64^. These treatments face limitations such as the possible emergence of drug-resistant mutants and side effects^65^, highlighting the need to identify new therapeutic approaches for timely control of the disease. Host-directed antiviral therapeutics, such as the TMPRSS2 inhibitor nafamostat^28,29^ and phosphoinositide 5-kinase inhibitor apilimod^30,31^, may offer an valuable alternative method to combat viral infections. Several host-directed antivirals against SARS-CoV-2 have been approved by FDA, or are in preclinical or clinical trials^66^, such as ER α-glucosidases I and II inhibitor (IHVR-1902) and co-translational translocation inhibitor that modulates SEC61 translocon in the ER (PS3061)^67^. With this rationale in mind, we investigated the antiviral efficacy of IMP-1088, an inhibitor of human NMT (NMT1 and NMT2), against SARS-CoV-2, RSV, SFV, and VSV infections. We found that IMP-1088 significantly inhibited the late stages of both SARS-CoV-2 and RSV infection cycles at nanomolar concentrations, whereases the infection of SFV and VSV was not affected, indicating that the trafficking, assembly and release mechanism of these enveloped viruses is different from that of SARS-CoV-2 and RSV.

Previous studies have also shown that inhibiting host NMT disrupts the infection of multiple human viruses, particularly those that contain *N*-myristoylated viral proteins, but not related enveloped viruses uses as controls^19–21^. The sequence of SARS-CoV-2 genome, however, does not harbour obvious *N*-myristoylation sites^68^. Yet, IMP-1088 demonstrated powerful inhibition of SARS-CoV-2 infection in various cellular models, including human lung and primary fully differentiated nasal epithelial cells, as well as human choroid plexus brain organoids. Interestingly, the nasal cells robustly express the entry factors ACE2 and TMPRSS2^26^, whereas the brain organoid model used in this study had no detectable levels of ACE2 and TMPRSS2. The antiviral activity of IMP-1088 extends to other SARS-CoV-2 variants, including Omicron and Delta, and to other viruses, including RSV. Together these results indicate that the anti-viral effect of IMP-1088 occurs through targeting of host functions, rather than viral functions, and that the drug is effective regardless of the viral entry route.

Although the IMP-1088 treatment did not interfere with SARS-CoV-2 entry, replication, or release, our results demonstrated that the SARS-CoV-2 virions released from IMP-1088-treated cells had significantly reduced levels of S proteins, which were also less cleaved by Furin, rendering the progeny virions less infectious. Our ultrastructural and live imaging studies showed that inhibition of host NMT causes an alteration of the secretory pathway, particularly from ER to ERGIC and Golgi, and eventually to plasma membrane. This seemed to activate an alternative SARS-CoV-2 egress pathway, that bypassed the ERGIC and Golgi transport and redirected the spike protein to lysosomal intermediate, disrupting the natural virion assembly and maturation pathways, and leading to a subsequent loss of the infectivity of released virions. Conversely, the infectivity of SFV and VSV, that are also enveloped viruses decorated by transmembrane highly glycosylated spikes which assemble at the host cell plasma membrane^69,70^, was not affected by the inhibition of host cell NMTs.

This raises the question of how IMP-1088 causes the disruption in the secretory pathway membrane trafficking. Our results demonstrated that IMP-1088 treatment led to significant alterations in the global proteome of A549-AT cells, particularly within the early secretory pathway and intracellular membrane trafficking. These findings align with previous studies indicating that protein *N*-myristoylation is essential for the sorting and trafficking of proteins to their appropriate cellular destinations, vesicular trafficking, membrane fusion, and signal transduction processes within the secretory pathway^71,72^. We also observed that loss of *N*-myristoylation led to a significant reduction in the abundance of proteins destined to be *N*-myristoylated, confirming earlier work where inhibition of NMT led to degradation of protein with a predicted *N*-myristoylation signal at their N-terminus^42^.

In vehicle control treated A549-AT cells, viral particles were typically observed within nuclear envelope, ER, ERGIC, Golgi, and lysosomes, while in cells treated with IMP-1088, viral particles were found in nuclear envelope, ER, and lysosomes but very rarely found in ERGIC or Golgi. Bypassing ERGIC and Golgi compartments evidently affects virion assembly and maturation, presumably by interfering with Furin cleavage, which normally occurs within the ERGIC and Golgi. Our studies with the S protein trafficking reveals that under this condition cells redirect immature transmembrane proteins from the ER to lysosomes and, unexpectedly, to the cell surface, providing a hitherto uncharacterized secretion mechanism that bypasses the Golgi. The physiological function of this pathway will require further studies.

Together, our findings underscore the prophylactic and therapeutic potential of NMT inhibitor IMP-1088 not only combating SARS-CoV-2 infection but also RSV, a leading cause of death in children. The loss of infectivity of the virions released from drug treated cells suggests that this antiviral strategy could potentially reduce the person-to-person transmission of the infection, which would have as significant impact on stalling the pandemic. This research paves the way for further exploration of the antiviral mechanisms and clinical applications of NMT inhibitors.

## MATERIALS AND METHODS

### A549-AT cells, drug treatments, and SARS-CoV-2

Human small lung carcinoma A549-AT cells stably expressing ACE2 and TMPRSS2-GFP^73^ and VeroE6^74^ cells were maintained in Dulbecco’s Modified Eagle’s High Glucose Medium (DMEM, Merck, Cat. no. D6546), 1% L-glutamine (Gibco, Cat. no. 35050061), 1% non-essential amino acids (NEAA, Gibco, Cat. no. 11140050), 10% foetal bovine serum (FBS, Thermo Fisher Scientific, Cat. no. 10270-106), 1x penicillin and streptomycin (P/S, Merck, Cat. no. P0781) and incubated at 37 °C in 5% CO_2_ atmosphere.

All A549-AT experiments analysed by imaging were performed in 96-well imaging plates (PerkinElmer, Cat. no. 6055302). Cells were seeded 24 h prior to infection at a density of 15.000 cells/well. The cells were treated with indicated concentrations of IMP-1088 (MedChemExpress, Cat. no. HY-112258), UCN-01 (Merck, Cat. no. 539644), or 4OH-Dynasore (Merck, Cat. no. D7693), at the indicated times before or after infection. The drug stocks (10 mM IMP-1088, 10 mM UCN-01, 100 mM 4OH-Dynasore) were prepared in DMSO (Merck, Cat. no. D2650), which was used in the experiments as vehicle control at a dilution correspondent to the amount of solvent contained in the highest concentration of each used drug. At indicated times after infection, cells were fixed and processed for indirect immunofluorescence detection of viral and cellular components (see Immunofluorescence staining below).

SARS-CoV-2 Wuhan (strain B.1), Delta (strain B.1.617.2) and Omicron (B.1.1.529) viruses were produced in Caco2 cells (passage 2)^74^ in the biosafety level 3 (BSL-3) facility of the University of Helsinki under permits 30 HUS/32/2018§16. Cells were infected in DMEM supplemented with 2% FBS, 2 mM glutamine, 1xP/S, 20 mM HEPES, pH 7.2, at MOI=0.01. Media containing viruses collected at 48 hpi. After two centrifugations at 300xg and 3000xg, for 5 and 10 min, respectively, the cleared media were aliquoted and stored at-80°C in the BSL3 premises. Viruses were tittered by standard plaque assay on VeroE6-TMPRSS2^74^. Deep sequencing confirmed that all viruses had an intact Furin cleavage site on the spike.

### Immunofluorescence staining

The cells in 96-well plates were fixed with 4% paraformaldehyde (PFA, Merck, Cat. no. P6148) in phosphate buffered saline (PBS), at room temperature for 20 min. PFA was then replaced with PBS and all the plates infected with SARS-CoV-2 were inactivated according to the institutional regulations using ultraviolet radiation (5000 J/m^2^ dose, UVP Crosslinker model CL-1000, Analytikjena) before transferring the samples out of the BSL-3 facility. The excess fixative was blocked by incubation with 50 mM ammonium chloride (NH_4_Cl; Merck Cat. no. 254134) prepared in PBS at room temperature for 20 min. Cells were then permeabilized with 0.1% (weight/volume) Triton X-100 (Merck, Cat. no. X100) in PBS and the nuclei were stained with the DNA dye Hoechst 33342 (1 mg/ml, Thermo Fisher Scientific, Cat. no. 62249) in Dulbecco-modified PBS (Merck, Cat. no. D8662) supplemented with 0.2% BSA (Merck, Cat. no. 9048-46-8), here referred to as Dulbecco-BSA, for 10 min. The cells were then washed twice with Dulbecco-BSA and incubated with the primary antibodies diluted in Dulbecco-BSA at +4°C, overnight. On the following day, the cells were washed twice with Dulbecco-BSA and incubated with the secondary antibodies diluted in Dulbecco-BSA for 1 hour at room temperature covered from light. After two washes in Dulbecco-BSA, cells were imaged in PBS. Viral nucleocapsid (N) protein was detected with an in-house-developed rabbit polyclonal antibody raised against the viral N protein of SARS-CoV-2 (a kind gift from Dr. Jussi Hepojoki^74^, University of Helsinki, Finland, dilution 1:2000). The mouse antibody against dsRNA (dilution 1:500) was from SCICONS (Cat. no. 10010500), Alexa Fluor 647-conjugated goat anti-rabbit and Alexa fluor 550-conjugated goat anti-mouse were from Thermo Fisher Scientific (Cat. no. A32733 and A32727, respectively). Actin was stained using Alexa Fluor-488 conjugated phalloidin (Invitrogen, Cat. no. A12379).

### Respiratory syncytial virus (RSV), Semliki Forest (SFV) and Vesicular stomatitis virus (VSV)

The three viruses have been previously engineered to express green fluorescent protein (GFP) as a reporter gene: SFV-GFP, VSV-GFP, RSV-GFP^35–37^. A549 cells, which are susceptible to the three viruses, were pre-treated with DMSO (vehicle control) or with 0.01, 0.1 or 1 µM IMP-1088 at 2 hbi or 1 hpi and infected with low doses (1:20 RSV for 24 h or 48 h; 1:200 VSV for 12 h or 24 h; and, ad SFV for 12 h or 24 h) of viruses. These infection periods account for early and late timepoints during respective viral infections, and the late infection timepoints were adjusted to allow multiple rounds of infection. Cells were fixed and stained with DAPI and imaged as explained below.

### High-throughput imaging and automated image analysis

Automated fluorescence imaging was done using a Molecular Devices Image-Xpress Nano high-content epifluorescence microscope equipped with a 10x and 20x objective and a 4.7-megapixel CMOS (complementary metaloxide semiconductor) camera (pixel size, 0.332μm). Image analysis was performed with CellProfiler-4 software (http://www.cellprofiler.org). Automated detection of nuclei was performed using the Otsu algorithm inbuilt in the software. To automatically identify infected cells, an area surrounding each nucleus (5-pixel expansion of the nuclear area) was used to estimate the mean fluorescence intensity of the viral N or dsRNA immunolabeling in each cell, using an intensity threshold such that <0.01% of positive cells were detected in noninfected wells. On average, >30,000 cells per sample were analysed. The percentage of cells positive for the virus N-protein, dsRNA or GFP-VSV, GFP-RSV, GFP-SFV relative to the total number of detected nuclei was then quantified and plotted.

### Human pseudostratified mucociliary nasal epithelial cell seeding and differentiation

All nasal epithelium experiment were conducted under the University of Queensland Ethical Approval 2022/HE002162. Human pseudostratified mucociliary nasal epithelial cells were seeded and differentiated as previously described^26^. In short, human nasal epithelial cells from donors were seeded at a density of 7.5 × 105 cells/trans-well on 6.5-mm trans-well polyester membranes with 0.4 μm pores (Corning Costar) and cultured in PneumaCult™-Ex Plus Medium (STEMCELL Technologies, Cat. no. 05041). Cells were monitored for confluence. When a confluent monolayer was achieved, cells were “air-lifted” by removing the media from the apical chamber and replacing the basolateral media with PneumaCult™-ALI-S Medium (ALI) (STEMCELL Technologies, Cat. no. 05050). Medium was replaced in the basal compartment 3 times a week, and the cells were maintained in ALI conditions for at least 3 weeks until ciliated cells and mucus were observed, and cells obtained a transepithelial electrical resistance (TEER) measurement greater than 800 Ω/cm2.

### Infection and immunostaining of ALI cells

The differentiated ALI cells were infected in a biosafety level 3 (BSL-3) laboratory and treated with DMSO (vehicle control), or 1 µM IMP-1088 for 2 h before infection, and then infected with MOI= 1*10^^6^ of SARS-CoV-2 for 72 h. Cells were fixed with 4% paraformaldehyde (Electron Microscopy Sciences, Cat. no. 15710) in PBS for 45 minutes at room temperature, followed by a blocking with 0.5% BSA (Sigma-Aldrich, Cat. no. A8022) in PBS for 30 minutes and permeabilization with 0.02% of Triton X-100 (Sigma-Aldrich, Cat. no. X100-100ML) in PBS for 15 minutes at room temperature. After washing twice with PBS/BSA and a second blocking step for 10 minutes at room temperature, samples were incubated with polyclonal rabbit anti-N protein of SARS-CoV-2 diluted in 0.5% BSA in PBS blocking solution: 1:2000 and incubated overnight at 4 °C. After 3 washing steps with 0.5% BSA/PBS for 5 minutes each time, the samples were incubated with secondary antibody (1:1000 dilution) Alexa Flour 647 donkey anti-rabbit (Invitrogen, Cat. no. A-31573) for 1 hour at room temperature in dark, and after 3 washes with 0.5% BSA/PBS, the cells. actin was stained using Alexa Fluor-488 conjugated phalloidin (Invitrogen, Cat. no. A12379) nuclei were stained with DAPI. The trans-well membranes with cells were cut with a scalpel, briefly dipped in milli-q water, and mounted on a class slide using ProLong Gold Antifade Mountant (Thermo Fisher Scientific, Cat. no. P10144). Mounted samples were imaged on a spinning disk confocal system (Marianas; 3I) consisting of an Axio Observer Z1 (Carl Zeiss) equipped with a CSU-W1 spinning disk head (Yokogawa Corporation of America), ORCA-Flash4.0 v2 sCMOS camera (Hamamatsu Photonics), and 63× 1.4 NA/Plan-Apochromat/180 μm WD objective.

### Paraffin sectioning and staining with H&E

ALI samples were collected and processed into paraffin using an automatic paraffin tissue processor TP1020. In brief the samples placed into a cassette and dehydrated with constant agitation in a graded series of ethanol, 50%, 70%, 95%, 100%, 100%, 100% ethanol, then cleared in 2 x Xylene 100% with vacuum for 40 min and then incubated in paraffin wax 90 mins with vacuum. Samples were blocked and the cut using a rotary microtome (Leica, RM2235), 10 µm thick section were stretched on a 40°C water bath and collected on positively coat slides (Uber). Paraffin sections on slides need to be dried down in the 60°C oven to remove water then dewaxed and rehydrated through a graded series of ethanol (2 × 100 % xylene for 10 min each and 2 × 100% ethanol 5 mins each, 70% ethanol for 3 min and then water 1 min.). ALI sections were stained with Mayer’s Hematoxylin (Sigma-Aldrich, Cat. no. 51275) for 2 mins, washed in water until the stain turned blue approximately 2 mins and then counter stained with 1% Eosin (Sigma-Aldrich, Cat. no. 17372-87-1) for 30 Sec, washed briefly in water and then dehydrated 70% ethanol, for 30 sec two changes of 100% ethanol and then incubated in Xylene before being coversliped in Ultramount permeant mounting medium.

### Human embryonic stem cells culture and cortical organoids generation

The human embryonic stem cells WTC iPSCs, which were a gift from Professor Bruce Conklin, were cultured in mTeSR (Stem Cell Technologies, Cat. no. 85851) on feeder-free hESCs medium on Matrigel (Stem Cell Technologies, Cat. no. 354277) according to the protocols detailed in (https://www.stemcell.com/maintenance-of-human-pluripotent-stem-cells-in-mtesr1.html).

### Choroid Plexus-Cortical organoids generation (ChPCOs)

The hPSC colonies were plated at a density of 20%–30% on a hESC-qualified basement membrane matrix on a 6-well plate to produce ChPCOs. One day before neuroectoderm (NEct) induction, hPSC colonies were kept alive with mTeSR. To produce NEct colonies, hPSC colonies were grown in N2 medium for three days: 1% MEM Non-Essential Amino Acids (Gibco, Cat. no. 11140-050), 1% penicillin/streptomycin (Gibco, Cat. no. 15140148), 0.1% β-mercaptoethanol (Gibco, Cat. no. 21985-023), 2% B-27 supplement (Gibco, Cat. no. 17504044), 1% N-2 supplement (Gibco, Cat. no. 17502-048), 1% MEM Non-Essential Amino Acids (Gibco, Cat. no. 11140-050), 1% penicillin/streptomycin (Gibco, Cat. no. 11320-33), and 0.1% β-mercaptoethanol (Gibco, Cat. no.11320-33). Every day, new N2 medium containing inhibitors was added. As we recently described^75^, on the fourth day, induced NEct colonies were elevated with dispase (2.4 unit/ml) to form neural spheroids. These spheroids were grown in N2 media for the following four days, with daily supplements of (bFGF, 40 ng/mL; R&D, Cat. no. 233-FB-01M), 2µM CHIR99021 (Sigma-Aldrich, Cat. no. SML1046-5MG), and 2ng/ml of BMP4 (Thermofisher, Cat. no. PHC9391). After being embedded in Matrigel, patterned neural spheroids were transferred to DMEM-F12, also known as Neurobasal medium (Gibco, Cat. no. A35829-01), the terminal differentiation medium. 17.5 µl of β-mercaptoethanol in 50 ml of medium (Sigma), 0.5% N2, 1% GlutaMAX, 1% MEM Non-Essential Amino Acids, 1% penicillin/streptomycin, 1% B-27 supplement, enhanced with 3µM CHIR99021 and 5ng/ml of BMP4. Three times a week, new media was added. The University of Queensland Human Research Ethics Committee gave its approval for all trials, which were conducted in compliance with its ethical principles (Approval number: 2019000159).

### SARS-CoV-2 infection of ChPCOs

SARS-CoV-2 isolate QLD1517/2021 (alpha variant, GISAID accession EPI_ISL_944644) passage 2 (P2) was kindly provided by the Queensland Health Forensic and Scientific Services, Queensland Department of Health. The isolate was amplified on Vero E6-TMPRSS2 cells to generate virus stock (P3) and sequenced confirmed^76^. Virus titers were determined by immuno-fluorescent foci-forming assay on Vero E6 cells^77^. Organoids were pre-incubated for 2 h in media containing DMSO or 1µM of IMP-1088 or 30 µM nafamostat, TMPRSS2 inhibitor, or 0.25 µM apilimod and infected with 10^6^ foci-forming units (FFU) of QLD1517 P3 for 6 h at 37 °C. After incubation, inoculum was removed, and organoids were washed 3 times before replacing with fresh media with corresponding drugs for 72 hpi before being processed for immunohistochemistry.

### Immunostaining and image quantification of ChPCOs

Tissue processing and immunostaining were carried out described previously^78^. Briefly, organoids were fixed in 4% PFA for 60 minutes at RT, then washed three times for 10 minutes at room temperature with 1x PBS. After that, fixed organoids were submerged in 30% sucrose in PBS at 4 °C and let to sink, then submerged in a solution consisting of 30% sucrose on dry ice and Optimal Cutting Temperature (O.C.T.) in a 3:1 ratio. Following serial slices at 14 µM thickness, mounted tissues were gathered onto Superfrost slides (Thermo Scientific, Cat. no. SF41296). Sectioned organoids were blocked for one hour with 3% BSA containing 0.1% triton X-100 and washed three times with PBS for 10 min at RT. Primary antibodies, polyclonal sheep anti-TTR (BIO RAD, Cat. no. AHP1837), polyclonal rabbit anti-N protein of SARS-CoV-2 (dilution 1:2000), guinea pig polyclonal anti-MAP-2 (synaptic system, Cat. no. 188 004, 1:1000) and rabbit monoclonal anti-cleaved caspase-3 (Asp175) (5A1E) (Cell Signalling, Cat. no. 9664) were incubated over night at 4°C, and then washed three times with PBS at RT for 10 minutes each. Before mounting and photographing, tissues were given an hour at RT incubation with the Alexa Fluor 647-conjugated goat anti-rabbit (Thermo Fisher Scientific, Cat. no. A32733), Alexa fluor 546-Goat anti-Guinea Pig IgG (H+L) (Thermo Fisher Scientific, Cat. no. A-11074), and Alexa fluor 488-conjugated Donkey anti-Sheep IgG (H+L) (Thermo Fisher Scientific, Cat. no. A-11015) secondary antibodies. The wholemount was carried out as prior to^79^. Hoechst 33342 (Invitrogen, Cat. no. H3570) was used to counterstained all the samples. All images were obtained using Zeiss AxioScan Z1. Image acquisition was optimized per immunofluorescent marker and captured with a z-stack of 4.5 μm, 0.8 μm intervals. Processed images were converted to TIFFs and nuclei were segmented as previously described^80^. Following nuclei segmentation, object detection for each marker was performed based on the maximum entropy, calculated by *S =-(sum)p*log2(p);* where p is the probability of a pixel greyscale value in the image^81^. Objects in each channel were stacked to generate composite images and multiplied to find the area and number of overlaid objects, for example number of cleaved caspase-3 positive nuclei, or infection positive MAP2 positive neurons. Density is calculated by the total area of a marker, normalized to the area of the organoid section. Image acquisition and quantification was performed in Zen 3.6, and ImageJ.

### Viral release and reinfection of ChPCOs

T175 flasks of Vero E6 cells were pre-treated with 1 µM IMP-1088 or DMSO (control) before infecting with 10^6^ foci-forming units (FFU) of QLD1517 P3. After 2 h, viral inoculum was removed, and the cells were washed 3 times before adding fresh media with the corresponding drugs. The virus was left to replicate for 3 days, and supernatant clarified and stored at – 80°C. RNA was harvested, and qRT-PCR done to normalize the viral copy number between DMSO-and IMP-1088 treated stocks. Following this, Vero E6 cells or organoids were infected with an equivalent viral copy number 1*10^6^ for 2 and 6 h, respectively. Cells and Organoid were washed 3 times with PBS and incubated in fresh media for 24 and 72 hpi, respectively. Samples were fixed for immunohistochemistry, imaging and quantification as explained above.

### scRNAseq of ChPCOs

Three choroid plexus (CP) organoids (60-day old) were pooled, washed once with DPBS and dissociated using Papain Dissociation Kit (Worthington Biochemical Corporation, Cat. no. LK003150) according to manufacturer’s instructions. Briefly, collected organoids were dissociated by incubation on a temperature-controlled shaker at 37°C at 190 rpm, and the enzymatic reaction was stopped by addition of stopping solution after 47 minutes. For quality control purposes, cell viability after dissociation was confirmed to be above 95% by Trypan Blue cell count and FACS analysis of DR (Thermo Fisher, Cat. no. D15106) positive cells. scRNAseq sample preparation was done using HIVE™ scRNAseq Solution kit (Perkin Elmer, Cat. no. NOVA-HCB018) by loading 30.000 live cells into each HIVE sample collector. In total, two HIVE sample collectors (containing pooled cells from 3 organoids) were used as technical replicates. Sample preparation and library preparation were done according to manufacturer’s instructions. Sequencing was run using NovaSeq 6000 SP Reagent Kit v1.5 (Illumina, Cat. no. 20028401) on a NovaSeq 6000 System (Illumina), estimating 36.000 reads/cell.

### scRNA-seq analysis

scRNA-seq data were aligned to the GRCh38.104 reference genome using BeeNet v1.1.X with the number of barcodes estimated at 40% of the total cells as indicated in the vignette. Double detection was performed to identify reliable singlets using DoubletFinder (v2.0.4)^82^ and quality control further removed cells with fewer than 500 genes per cell as well as cells with greater than 20% mitochondrial content. As a result, 6952 high-quality cells were retained for downstream analyses. To perform dimensionality reduction and clustering, we processed the data with Seurat v5^83^. Raw data were normalised via the LogNormalize method with scale.factor of 10,000 using the NormalizeData function. Highly variable genes were identified using the variance stabilizing transformation method with default parameters. Based on these genes, a PCA was performed and the first 30 PCs were retained for clustering and visualization via UMAP. Cell clusters were identified via the ‘FindClusters’ function via the Louvain algorithm with a resolution parameter of 0.6, resulting in 14 clusters. Cluster-specific genes were identified via the Find All Markers function via the Wilcoxon test and used to assign cell-type labels. Markers were selected with logfc. threshold at 0.25 and minimum percentage of expression in 5% of total cells. Cluster markers were interpreted and assigned cell type identity by ScType^84^ and known literature cell type annotation. Choroid plexus (TTR and AQP1), Cortical hem (MSX1), Ependymal (MCIDAS^85^), Radial glia (SOX2, PAX6 and HOPX), Astro-progenitor (SLC1A3), Excitatory neurons (SLC17A6, MAP2 and FOXP2) and Inhibitory neurons (GABRA2, MAP2 and RELN) were annotated with their canonical markers (Supplementary Figure 2). To visualize the marker expression and COVID-related genes, scCustomize v2.0.1^86^ was applied to generate the dot plots and violin plots.

### qRT-PCR

Viral RNA was extracted from 50 ml cell culture medium using QIAamp Viral RNA Mini Kit according to protocol provided by the manufacturer (Qiagen; Cat. no. 52904). RNA concentration and quality were evaluated using NanoDrop 2000 Spectrophotometer (Thermo Fischer Scientific). The qRT-PCR was run in quadruplicate using TaqMan Fast Virus 1-step MasterMix (Thermo Fischer Scientific, Cat. no. 4444432). Primers and probes used for the qRT-PRC reactions were ordered from Metabion international AG (www.metabion.com) as follows: RdRP-SARSr-F2: 5’-GTG ARA TGG TCA TGT GTG GCG G-3’ forward primer, RdRP-SARSr-R2: 5’-CAR ATG TTA AAS ACA CTA TTA GCA TA – 3’ reverse primer, RdRP-SARSr-P2: 5’-6-Fam-CAG GTG GAA CCT CAT CAG GAG ATG C-BHQ-1-3’ the probe. The qRT-PCR reaction was run with Agilent Technologies Stratagene Mx3005P using the following parameters: Reverse transcription for 5 min at 50°C, initial denaturation for 20 s at 95°C and amplification steps at 95°C for 3 s and 60°C for 30 s (the amplifications steps were repeated for 40 cycles).

### Sequencing of virus isolates

To test if the inhibition of NMT-1 and 2 induced mutations in the released SARS-CoV-2 progeny virion in A549-AT treated with IMP-1088 or DMSO control, viral RNA was extracted from 50 ml cell culture medium using QIAamp Viral RNA Mini Kit according to protocol provided by the manufacturer (Qiagen, Cat. no. 52904). RNA concentration and quality were evaluated using NanoDrop 2000 Spectrophotometer (Thermo Fischer Scientific) and reverse-transcribed to cDNA using LunaScript® RT SuperMix Kit (New England Biolabs, Cat no. E3010) with random hexamers and oligo dT. SARS-CoV-2-specific amplicons were generated using xGen Artic V4 NCoV-2019 primers (Integrated DNA Technologies Inc., Coralville, Iowa) and Q5 Hot Start High Fidelity PCR Master Mix (New England Biolabs, Cat. no. E0555S). Sequencing libraries were prepared using NEBNext ultra II DNA library kit (New England Biolabs, Cat. no. E7805S) and unique dual indexes (Integrated DNA Technologies Inc.) according to the manufacturer’s instructions, followed by sequencing using Illumina Miseq with v3 sequencing kit. The virus genomes were assembled using HaVoC-pipeline^87^, using fastp^88^ for trimming, BWA-MEM for assembly^89^, LoFreq version 2^90^ for single nucleotide variant calling and BCFTools^91^ for consensus sequence calling.

### SARS-CoV-2 purification

VeroE6 were grown to 70-80% confluency in growth media and treated with 1 µM of IMP-1088 or DMSO for 2 hbi and then infected with SARS-CoV-2 Wuh (MOI=1) for 2 h. Cells were washed 3 times with PBS and new growth media was added containing 1 µM IMP-1088 or DMSO for 72 hpi. First step is the media clearance by centrifuging the media containing released virus twice for 20 min, 10,000× g at 4°C. Second step is SARS-CoV-2 Pelleting The pre-cleared virus-containing media was layered on top of a 5 mL 30% (w/v) sucrose cushion in buffer containing 20 mM HEPES and 155 mM NaCl at pH 7.0 (HN) and centrifuged for 1.5 h at 100,000 × g at 4°C. the resulted Viral pellet was suspended in 200 µL PBS. Third step is SARS-CoV-2 purification. The pelleted virus was layered on top of sucrose gradient starting from the bottom to top, 60%-30%-255-12.5% (w/v) sucrose cushion in buffer containing 20 mM HEPES and 155 mM NaCl at pH 7.0 (HN) and centrifuged for 1.5 h at 100,000× g at 4°C. The results cloudy layer between 30% and 60% layers was collected as purified virus.

### Western blot

The purified virions were quantified using qRT-PCR as described above. 10*10^6^ virions of IMP-1088 or DMSO released purified viruses were lysed in 4x Laemmli protein sample buffer (BIO-RAD, Cat. no. 1610747). Samples were electrophoresed on 4-20% precast polyacrylamide gel (BIO-RAD, Cat. no. 4561093EDU) for 1 hour with 100 fixed voltage and transferred to PVDF (Bio-Rad, Cat.no. 1620177) membranes using wet method for 90 minutes with100 fixed voltage. Membranes were then blocked with 5 % Skim Milk for 1 hour at room temperature. Membranes were then incubated with the overnight at 4°C, with rabbit polyclonal anti-SARS-CoV-2 (2019-nCoV) Spike RBD Antibody (Sino Biological Inc, Cat. no. 40592-T62, 1: 1000) and rabbit polyclonal anti-SARS-CoV-2 (2019-nCoV) Spike S2 Antibody (Sino Biological Inc, Cat. no. 40590-T62, 1: 1000) or N protein of SARS-CoV-2 (dilution 1:2000) or SARS-CoV-2 membrane ab (R&D Systems, Cat no. MAB10690). Followed 3 time washing with PBST (PBS with 0.1 tween 20) and then incubated 1 hr with secondary antibodies IRDye 680LT-donkey anti-mouse IgG (LI-Cor, Cat. no. 926-68022) or IRDye 800CW-donkey anti-Rabbit IgG (LI-Cor, Cat. no. 926-32213) diluted in 0.1 BSA (1:10000) followed by 3 times PBST washing. Then, proteins were then visualized by ECL detection solution using LI-COR system.

### Proteomics analysis using high resolution Orbitrap mass spectrometry

The LC/MS/MS identification and quantification of the digested peptides of A549 cells treated with DMSO (Control) or 1 µM IMP-1088 for 48 h were performed using a S-TrapTM (Protifi, NY) Micro Spin Column Digestion protocol. Briefly, samples were solubilised by adding 50 µL of S-Trap lysis buffer (10% sodium dodecyl sulphate (SDS) in 100 mM Tris, pH 8.0) to 50 µL of sample, before reducing by adding 20 mM of dithiothreitol (DTT) and heating at 70°C for 60 min. Cysteine residues were alkylated to prevent disulphide bond reformation using 40 mM iodoacetamide for 30 min at room temperature in the dark. 2.5 µL of 12% phosphoric acid was added, followed by 165 µL of S-Trap binding buffer (90% methanol in 100 mM Tris) to the acidified lysate. The sample mix was then centrifuged through the S-Trap column at 4,000 x g for 1 min followed by three washes with 150 µL S-Trap binding buffer, with 4,000 x g centrifugation between each wash. Peptide digestion was initiated by adding 25 µL of 50 mM ammonium bicarbonate buffer (pH 8) containing 2 µg trypsin (Sequencing Grade Modified Trypsin, Promega, Cat. no. V5117) directly on top of the column and incubating overnight at 37°C. Peptides were eluted by three successive aliquots of 40 µL of 5%, 50%, and 75% acetonitrile in 0.1% formic acid, respectively. Eluted peptides were dried down using a vacuum concentrator (Concentrator Plus, Eppendorf). Samples were redissolved in 20 µL of 5% ACN (aq) and 2 µl were injected to a trap column (Thermo 22 mm x 300 µm, 5 µm, C18) at a flow rate of 10µL/min. Following 3 min wash the trap column was switched in-line with a resolving column (Water nanoEase 100 mm x 150 µm, 1.8 µm, 100 Å). The samples were eluted by a gradient was held constant at 8% for 4 min, then was increased linearly to 24% at 47 min, to 40% at 53 min and to 95% at 57 min. The gradient held constant for 1 min, before returning to start condition at 8% over 1 min. Mass spectrometry using LC-MS/MS was performed using Ultimate UHPLC system coupled to an Exploris 480 mass spectrometer with a FAIMS Pro interface (Thermo Fisher Scientific) The FAIMS compensation voltages were-45 and-65 V. The electrospray voltage was 2.2 kV in positive-ion mode, and the ion transfer tube temperature was 295°C. Full MS scans were acquired in the Orbitrap mass analyser over the range of m/z 340-1110 with a mass resolution of 120,000. The automatic gain control (AGC) target value was set at ‘Standard’, and the maximum accumulation time was ‘Auto’ for the MS. The MS/MS ions were measured in 6 windows from mass 350-470, in 18 windows from mass 465-645 and 5 windows from mass 640-1100 with an overlap of 1 m/z and quadrupole isolation mode. Analysis of data were performed using Spectronaut against a reference proteome with a Q-value cut-off of 0.05.

### Additional processing and statistical analysis of label-free mass-spectrometry

The standard, non-normalised output of Spectronaut (BGS Factory Report text file) was imported into R version 4.05 for further processing using a modified method described previously^92^. For proteins that mapped to multiple annotations, the ‘best annotation’ method previously described was used^92^. Each sample group was permitted to have up to one missing value (that is, an intensity not reported one or three individual replicates). Protein intensities of the parent group protein ‘PG.ProteinGroups’ were then log2 transformed and globally normalised using the quantile normalisation method^93^. Missing values were assumed to be missing at random and imputed using the knn nearest neighbour averaging method^94^. Following, unwanted sources of technical variation were removed by surrogate variable analysis ^95^. Sample clustering was confirmed using unsupervised principal component analysis and hierarchical clustering. Principle Component Analysis of the first two principal components using singular value decomposition, as implemented in pcaMethods version 1.82.0 and hierarchical clustering using ‘ward’ method for clustering distance ‘euclidean’. For hierarchical clustering, probabilities of clustering were determining using 100 bootstrap replications, as implemented in pvcluster version 2.2-0, and probabilities of the branch positions shown in the plot, together with the probabilities of clusters (red boxes). For differential protein abundance, generalised linear modelling with Bayes shrinkage as implemented in limma version 3.46.0 was performed and proteins were considered differentially abundant at a corrected p-value of 0.05 unless specified otherwise (adjusted for false discovery rate using the Benjamini and Hochberg method)^96^.

### Gene Set Enrichment Analyses

Heatmaps for proteins belonging to a gene set (gene ontology term) were independently generated by extracting all Gene Symbols from ‘org.Mm.eg.db’ version 3.1.2 that matched a specific GO term of interest. This was then filtered by proteins that were differentially abundant between IMP-1088 treated A549 cells versus DMSO treated A549 cells. The z-score of the protein intensities across all samples (row-wise) was then calculated across samples and plot as a heatmap using the pheatmap version 1.0.12. Clustering distance was ‘euclidean’ and clustering method ‘complete’ linkage. Targeted pathways selected included: ‘Mitochondria (GO:0005739)’, ‘Presynapse (GO:0098793)’, ‘the endoplasmic reticulum (GO:0005783)’, ‘endoplasmic reticulum–Golgi intermediate compartment (GO:0005793)’, ‘ER to Golgi transport vesicle (GO:0030134)’, ‘Golgi apparatus (GO:0005794)’, ‘secretory granule (GO:0030141)’, ‘transport vesicle (GO:0030133)’, and ‘ribosome (GO:0005840)’.

### Cell viability assay

To determine the effects of IMP-1088 on A549 cell viability, A549 cells were seeded in 96-well plates and grown to 70% confluence and treated with DMSO as a control, 10 µM UNC-01, 5 µM Puromycin, (0.01, 0.1, 1 µM) of IMP-1088 for 24 and 48 h. Cell viability was assessed using CellTiter-Glo® Luminescent Cell Viability Assay (Promega, Cat. no. G7570) as per manufacturer’s instructions.

### Electron microscopy (EM) analysis

Transmission electron microscopy analysis was performed on cultured A549-AT treated with DMSO vehicle (control) or 1 µM IMP-1088 for 48h and (± SARS-CoV-2 wuh). For structural analysis cells were fixed with 1.5% glutaraldehyde (Electron Microscopy Sciences, Cat. no. 16210) and 1.5% PFA in 0.1 M sodium cacodylate (Sigma-Aldrich, Cat. no. C0250) buffer pH 7.4 for 20 min at room temperature, washed three time for 3 min with 0.1 M sodium cacodylate buffer, and processed for EM using standard protocols. All samples were contrasted with 1% osmium tetroxide and 2% uranyl acetate before dehydration and embedded in LX-112 resin using BioWave tissue processing system (Pelco) as previously described^97^. Thin sections (80-90 nm) were cut using an ultramicrotome (Leica Biosystems, UC6FCS) and were imaged with a transmission electron microscope (JEOL USA, Inc. model 1101) equipped with cooled charge-coupled device camera (Olympus; Morada CCD Camera).

### Electron tomography (ET) 3D model

A549-AT cells were treated with 1 µM IMP1-088 1 hbi and infected with SARS-CoV-2 wuh at MOI=5 for 48 h and processed for EM as described above. Following embedding, approximately 200 nm resin sections were cut on a Leica Ultracut 6 ultramicrotome, and the grid was subsequently coated with a thin carbon layer. Tomography was completed as described previously^98^. In brief, a dual-axis tilt series spanning ± 60° with 1° increments was acquired on a Tecnai F30 transmission electron microscope (FEI Corp.) operating at 300 kV. Images were collected with a 4k × 4k OneView CMOS camera (Gatan Corp.) at 3,900× magnification, providing a pixel size of 30 Å. The specimens were tilted at 1-degree increments using a high tilt specimen holder between +60° and ±70°. Automated acquisition of the tilt series was carried out using the tomography software package SerialEM software^99^. Tilt series were reconstructed using weighted back-projection and patch tracking in IMOD (https://bio3d.colorado.edu/imod/). Segmentation and tomographic reconstructions were analysed and modelled with Amira (TGS Inc.).

### Immuno-EM

Immuno-EM analysis was performed on cultured A549-AT treated with DMSO vehicle (control) or 1 µM IMP-1088 for 30 min and infected with SARS-CoV-2 wuh at MOI=5. At 2 hpi, infection media was replaced with fresh culture media containing either DMSO vehicle (control) or 1 µM IMP-1088, and the cells were infected for 48 h. Immuno-EM sample preparation was done as follows: Cells were fixed with paraformaldehyde-lysine-periodate-fixative for 2 h^100^. Cells were permeabilized with 0.01% saponin (Sigma-Aldrich), 0.1% BSA in 100 mM Sodium Phosphate buffer pH 7.4 and immunolabelled with anti-calreticulin (Abcam, Cat. no. ab22683), anti-ERGIC-53 (Proteintech, Cat. no. 13364-1-AP) or anti-GM130 (BD Biosciences Cat. no. 610822) followed with 1.4 nm nanogold-conjugated anti-mouse (Nanoprobes, Cat. no. 2001) or anti-rabbit (Nanoprobes, Cat. no. 2003) secondary antibodies. Signal was gold enhanced using Gold Enhance EM Kit (Nanoprobes, Cat. no. 2113). Finally, cells were processed for osmication, dehydration, Epon embedding using BioWave tissue processing system, followed by sectioning and imaging as described above.

### Creation of pS^wt^ construct

The synthetic gene *nCoV-Fin-S-pEBB* was assembled from synthetic oligonucleotides and/or PCR products. The fragment was inserted into pEBBHA-N_AE812. The plasmid DNA was purified from transformed bacteria and concentration determined by UV spectroscopy. The final construct was verified by sequencing. The sequence identity within the insertion sites was 100%. The vector plasmid is pEBB-N-HA (N-terminal HA-tag) and the gene was first synthesized by GeneArt and then subcloned into a BamH1 site in pEBB-N-HA.

### Transient transfections

To study the effects of IMP-1088 on early secretory pathway structure in live cells, GFP-ERGIC53, ARF1-GFP plasmids were used. Briefly, A549 cells were transfected with 2 µg DNA of ARF1-GFP^48^ (ARF1-GFP was a gift from Paul Melancon, Addgene plasmid #39554; http://n2t.net/addgene:39554; RRID:Addgene_39554), Hsp47-GFP-KDEL^51^ (a gift from Eija Jokitalo, Univeristy of Helsinki, Finland) or pS^wt^ using Lipofectamine LTX and Plus Reagent (Invitrogen, Cat. no. 15338-100) as per manufacturer’s instructions. Cells were incubated in transfection media overnight at 37°C, 5% CO_2_, and then treated either with 1 µM IMP-1088 or 5 µg/ml of Brefeldin A. Cells were imaged using a Zeiss Plan Apochromat 63x/1.4 NA oil-immersion objective on a confocal/two-photon laser-scanning microscope (LSM 710; Carl Zeiss Pty Ltd, Australia) built around an Axio Observer Z1 body (Carl Zeiss) and equipped with two internal gallium arsenide phosphide (GaAsP) photomultiplier tubes (PMTs) and three normal PMTs for epi-(descanned) detection and two external GaAsP PMTs for non-descanned detection in two-photon imaging, and controlled by Zeiss Zen Black software. Movies were acquired at 2 frames per second (50-100 frames total) and analysed using Fiji2/Image J2. The image analysis protocol described below is modified from^101^.

### Immunostaining of GM130 and ERGIC-53 with SARS-CoV-2 S protein

To study how IMP-1088 impact the early secretory pathway structure, A549-AT cells were treated with 1 µM of IMP-1088 or DMSO (control) and (±SARS-CoV-2, MOI=1). 48 hpi, cells were cells fixed with 4% paraformaldehyde (ProSciTech, Cat. no. C004) in PBS for 30 min at RT, permeabilized with 0.1% of Triton X-100 (Sigma-Aldrich, Cat. no. X100-00ML) in PBS for 5 min (RT), blocked with 2% Bovine Serum Albumin (BSA) (Sigma-Aldrich, Cat. no. A8022-10G) in PBS (BSA/PBA) for 45 mins (RT), and then stained against mouse monoclonal primary anti-GM130 (BD Biosciences, Cat. no. 610822, 1:500) or rabbit polyclonal primary anti-ERGIC-53 (Proteintech, Cat. no. 13364-1-AP, 1:500) with or without rabbit polyclonal anti-SARS-CoV-2 (2019-nCoV) Spike RBD Antibody (Sino Biological Inc, Cat no. 40592-T62) and rabbit polyclonal anti-SARS-CoV-2 (2019-nCoV) Spike S2 Antibody (Sino Biological Inc, Cat. no. 40590-T62) were incubated over night at 4°C, and then washed three times with PBS at RT for 5 minutes each. Cells were then incubated with Alexa Fluor 647-conjugated goat anti-rabbit (Thermo Fisher Scientific, Cat. no. A32733), and Alexa fluor 550-conjugated Goat anti-Mouse (Thermo Fisher Scientific, Cat. no. A32727) secondary antibodies. Hoechst 33342 (Invitrogen, Cat. no. H3570) was used for nuclear stain and actin was stained using Alexa Fluor-488 conjugated phalloidin (Invitrogen, Cat. no. A12379) for 1 hour in dark at RT. Samples were washed 3 times with PBS and mounted using Prolong Diamond (Life Technologies Australia Pty Ltd, Cat no. P363961). Samples were imaged on a spinning-disk confocal system (Marianas; 3I, Inc.) consisting of a Axio Observer Z1 (Carl Zeiss) equipped with a CSU-W1 spinning-disk head (Yokogawa Corporation of America), ORCA-Flash4.0 v2 sCMOS camera (Hamamatsu Photonics), using 63x 1.2 NA C-Apo oil objective. Image acquisition was performed using SlideBook 6.0 (3I Inc) and processed using Fiji2/Image J2 (v2.14.0/1.54f).

Colocalization analysis was performed per cell on Max projected Z stacks by running Fiji plugin ‘coloc 2’ [https://github.com/fiji/Colocalisation_Analysis]. The spot detection protocol was done as previously described^101^ with the following modifications. After max projection, images were converted to Greyscale and threshold adjusted to exclude background. Following, the image was ‘smooth’-ed before creating a binary using existing Fiji plugins. After processing, the ‘Analyse Particles’ function was run with size (0-infinity pixel^2), circulatory (0.00-1.00) and edges excluded. The following parameters were measured: Cell and spot area (size, µm^2^) and number of spots. Minimum fluorescence (Min), and Maximum fluorescence (Max). The output files are exported to Excel for processing and collating, then graphs and statistical analysis was performed using Graph Prism 9.

### Live cell dynamics of ARF1 using confocal microscopy

A549-AT cells were transiently transfected with ARF1-GFP and treated with pharmacological inhibitors, IMP-1088 (1 μM) or Brefeldin-A (5 μg/ml) at time of the imaging. For paired analysis, the regions of interest were imaged at t_0 s_ (untreated control), and at t_40 s_ and t_240 s_ after drug treatment. Due to slow and cumulative effects of IMP-1088 on cells, different regions of interest were images at t_0 s_ (DMSO control), and at t_1 h_ and t_48_ h after drug treatment. Cells were imaged at a single optical plane, 2 frames per second, for 100-500 frames using a Zeiss Plan Apochromat 63x/1.4 NA oil-immersion objective on a confocal/two-photon laser-scanning microscope (LSM 710; Carl Zeiss Pty Ltd, Australia) built around an Axio Observer Z1 body (Carl Zeiss) and equipped with two internal gallium arsenide phosphide (GaAsP) photomultiplier tubes (PMTs) and three normal PMTs for epi-(descanned) detection and two external GaAsP PMTs for non-descanned detection in two-photon imaging, and controlled by Zeiss Zen Black software. Imaging was done at 37°C and in 5% CO_2_ atmosphere. Colocalization and spot detection analysis were performed per cell, as described above using Fiji2/Image J2.

### Immunostaining of internal and secreted proteins

To study how IMP-1088 impact the intracellular pS^wt^ trafficking, A549-AT cells were transfected with 2 µg DNA of pS^wt^ using Lipofectamine LTX and Plus Reagent (Invitrogen, Cat. no. 15338-100) as per manufacturer’s instructions. Cells were incubated in transfection media overnight at 37°C, 5% CO_2_, and then treated either with IMP-1088 (1µM for 48 h), Brefeldin A (5 µg/ml for 25 min) or DMSO (control, 48 h). Cells were cells fixed with 4% paraformaldehyde (ProSciTech, Cat. no. C004) in PBS for 30 min at RT and blocked with 2% BSA (Sigma-Aldrich, Cat. no. A8022-10G) in PBS (BSA/PBA) for 45 mins (RT) and then stained against external calreticulin using mouse monoclonal anti-calreticulin (Abcam, Cat. No. ab22683, 1:500), or mouse monoclonal primary anti-LAMP1 (Cell Signalling, Cat. no.15665, 1:500) or rabbit polyclonal anti-SARS-CoV-2 (2019-nCoV) Spike S2 Antibody (Sino Biological Inc, Cat. No. 40590-T62, 1:1000) at 4 °C overnight. Cells were then wash 3 times with PBS and stained with Alexa fluor 550-conjugated goat anti-mouse (Thermo Fisher Scientific, Cat. no. A32727, 1:1000) or Alexa fluor 488-conjugated goat anti-rabbit (Thermo Fisher Scientific, Cat. no. A-110081:1000) secondary antibodies for 1 h (at RT) followed by three washes with PBS. To immunolabel the intracellular calreticulin, Lamp-1 and pS^wt^, cells then fixed for 5 min with 4% PFA followed by 3 times of PBS wash and then permeabilized with 0.1% of Triton X-100 (Sigma-Aldrich, Cat. no. X100-00ML) in PBS for 5 min at RT, blocked with 2% BSA in PBS (BSA/PBA) for 45 mins at RT, and then stained against mouse monoclonal primary anti-GM130 (BD Biosciences, Cat. no. 610822, 1:500), anti-calreticulin, anti-LAMP1 and anti-SARS-CoV-2 Spike S2 antibodies as described above at 4°C overnight. Cells were washed 3 times with PBS and stained with Alexa Fluor 647 conjugated Goat anti-Mouse IgG (H+L) (Thermo Fisher Scientific, Cat. no. A-21236, 1:1000) and Alexa fluor 488-conjugated Goat anti-Rabbit (Thermo Fisher Scientific, Cat. no. A-11008, 1:1000) Secondary Antibodies with DAPI for the nuclear stain, and them embedded in ProLong Gold Antifade Mountant (Thermo Fischer Scientific, Cat. no. P36930).

### Furin rescue of released virions

To investigate Furin rescue of the spike cleavage of IMP-1088 released virions would increase their infectivity, 10^10^6^ of DMSO or IMP-1088 released virions were treated with two-fold dilutions of Furin (New England Biolabs, Cat. no. P8077L) in 20 mM HEPES, 0.1% Triton X-100, 1 mM CaCl_2_, 0.2mM β-mercaptoethanol (pH 7.5 at 25°C) in a 25 µl reaction. The reaction mix was incubated at 25°C for 6 hours. Then we tested the infectivity of those virions and Furin no treated DMSO or IMP-1088 released virions on VeroE6 cells for 24 h and then fixed and processed for immune staining of N protein of SARS-CoV-2 and DAPI and imaged and quantified as described in the above section.

### Statistical analysis

Statistical tests were conducted in GraphPad Prism 9 for macOS version 9.1.1. No statistical methods were used to predetermine sample sizes. The normality of the data was tested with Kolmogorov–Smirnov tests, and nonparametric tests and parametric tests were used to compare two independent groups (Mann–Whitney test or t test) and multiple groups (Kruskal– Wallis test or ordinary 2way ANOVA with Tukey post hoc multiple comparison test). Dot plots indicate average ± SEM, and violin plots show median ± quartiles together with data points from individual acquisitions or ROIs unless otherwise stated. The samples/cells were randomized for examination. Sex was not considered. The investigators were not blinded during experiments. Specific statistical tests for each experiment are described in the figure legends. P values of less than 0.05 were considered significant. Measurements were taken from distinct samples.

---

**Supplementary Figure 1.**
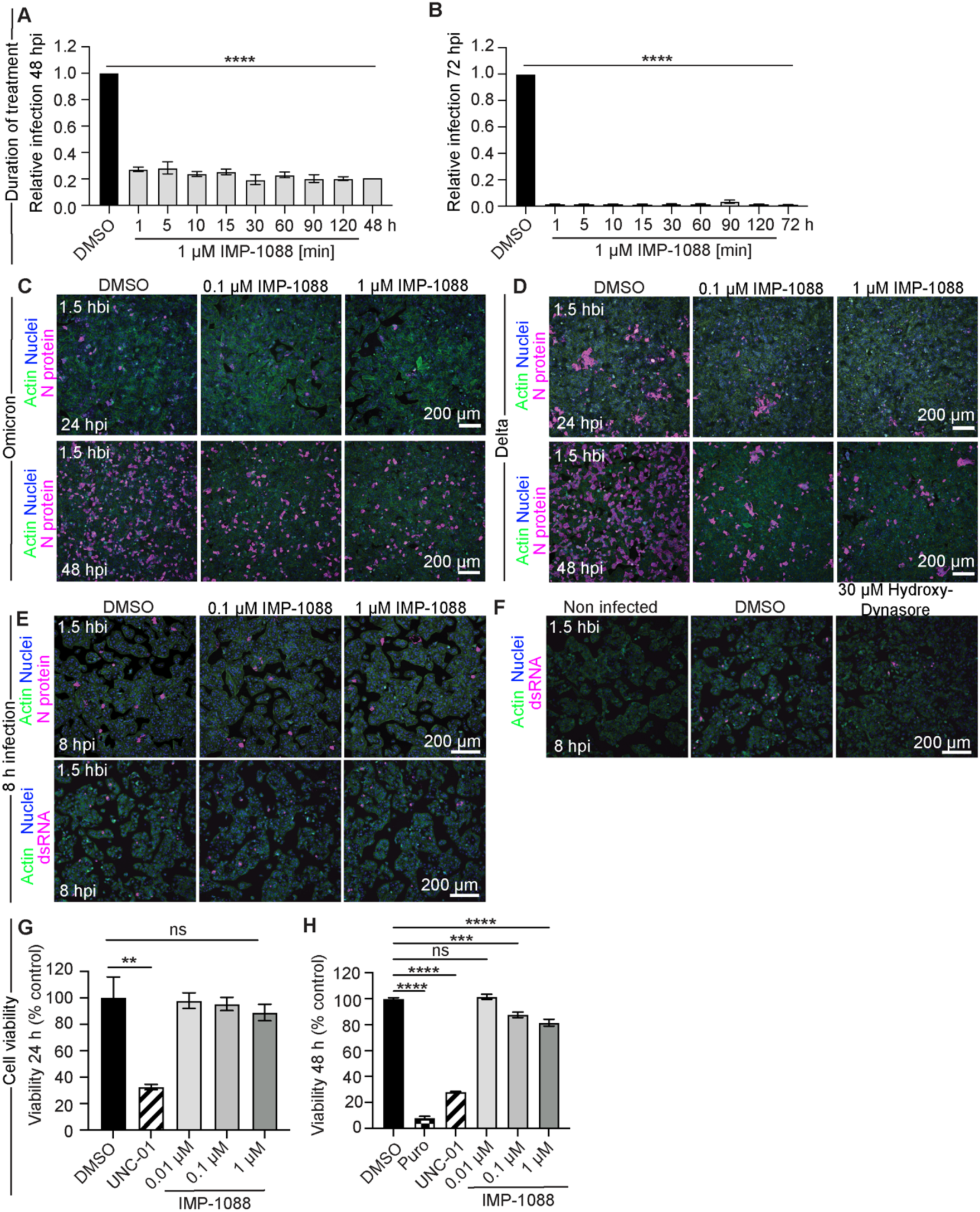
IMP-1088 does not impact viral entry, exhibits minimal cytotoxicity, and irreversibly inhibits NMT1 and-2 in a very short time. (A, B) Quantification of relative SARS-CoV-2 Wuh infections in A549-AT cells treated with DMSO or 1 µM IMP-1088 for indicated treatment times at 48 hpi (A) or 72 hpi (B). (C, D) Representative immunofluorescence images of A549-AT cells treated with indicated concentrations of IMP-1088 or DMSO vehicle control at 1.5 hbi SARS-CoV-2 Omicron (C) or Delta (D) infection for 24 h (top panels) and 48 h (bottom panels), respectively. Immunostainings show viral markers N (magenta), actin (green) and nuclei (blue). (E, F) Representative immunofluorescence images of A549-AT cells treated with indicated concentrations of IMP-1088 or DMSO vehicle control (E) or 30 µM hydroxy-dynasore (F) at 1.5 hbi of SARS-CoV-2 Wuh for 8 h. Immunostainings show viral markers N and dsRNA (magenta) actin (green) and nuclei (blue). (G, H) Cell viability quantification of A549-AT cells treated with indicated concentrations of IMP-1088, DMSO, 10 µM UNC-01, or 2.5 µM puromycin for 24 h (G) or 48 h (H). All the results are expressed as mean ± SEM of 3 biological replicates in each condition. Statistical analysis was performed with ordinary one-way ANOVA multiple comparisons test, **P<0.01, ***P<0.001 ****P<0.0001, ns=non-significant.

**Supplementary Figure 2.**
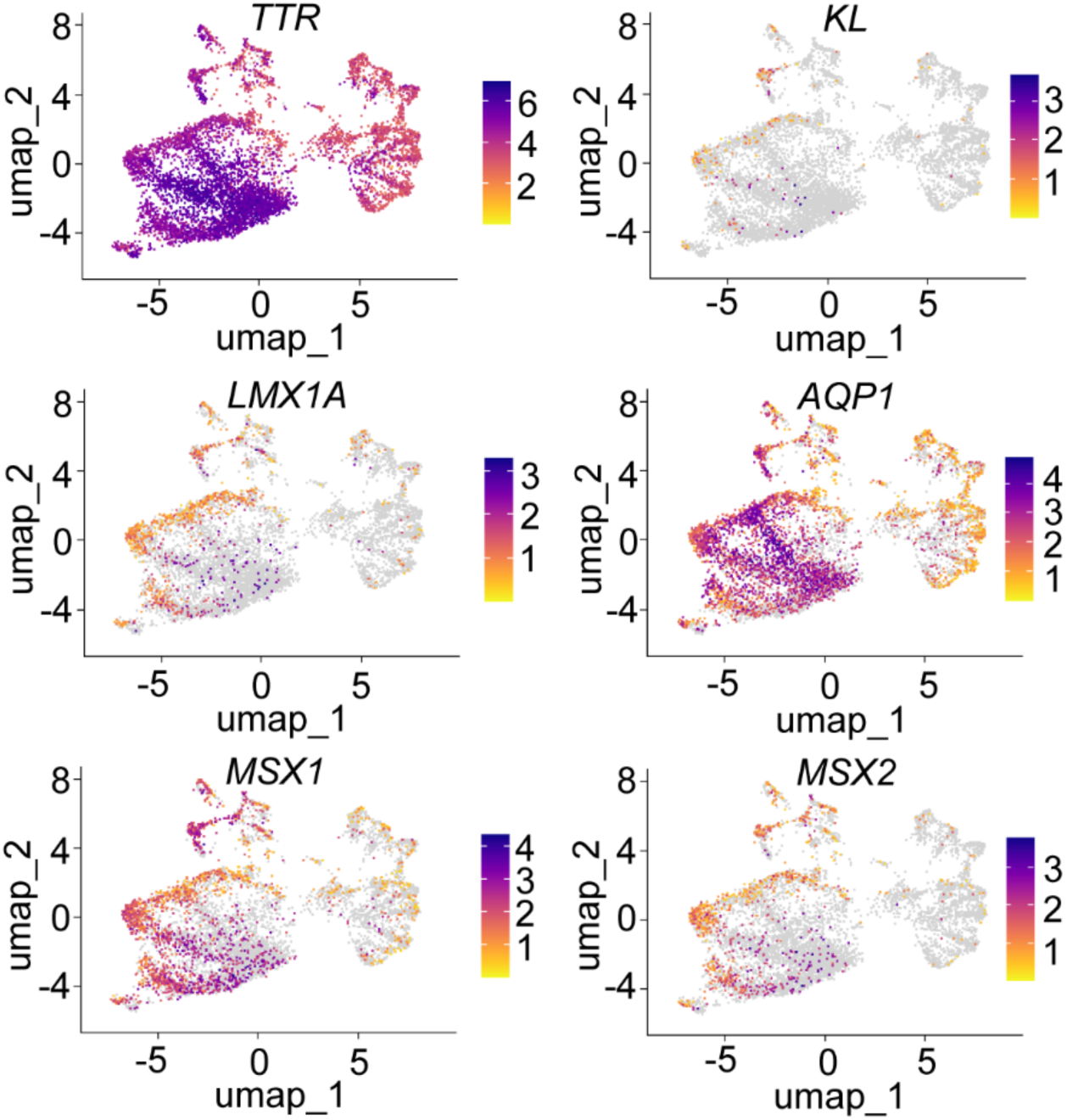
Marker gene expression visualisation from scRNA-seq data. Collection of marker expression for choroid plexus cells in CPCOs visualised in UMAP. Colour scale represents log transformed counts across cells.

**Supplementary Figure 3.**
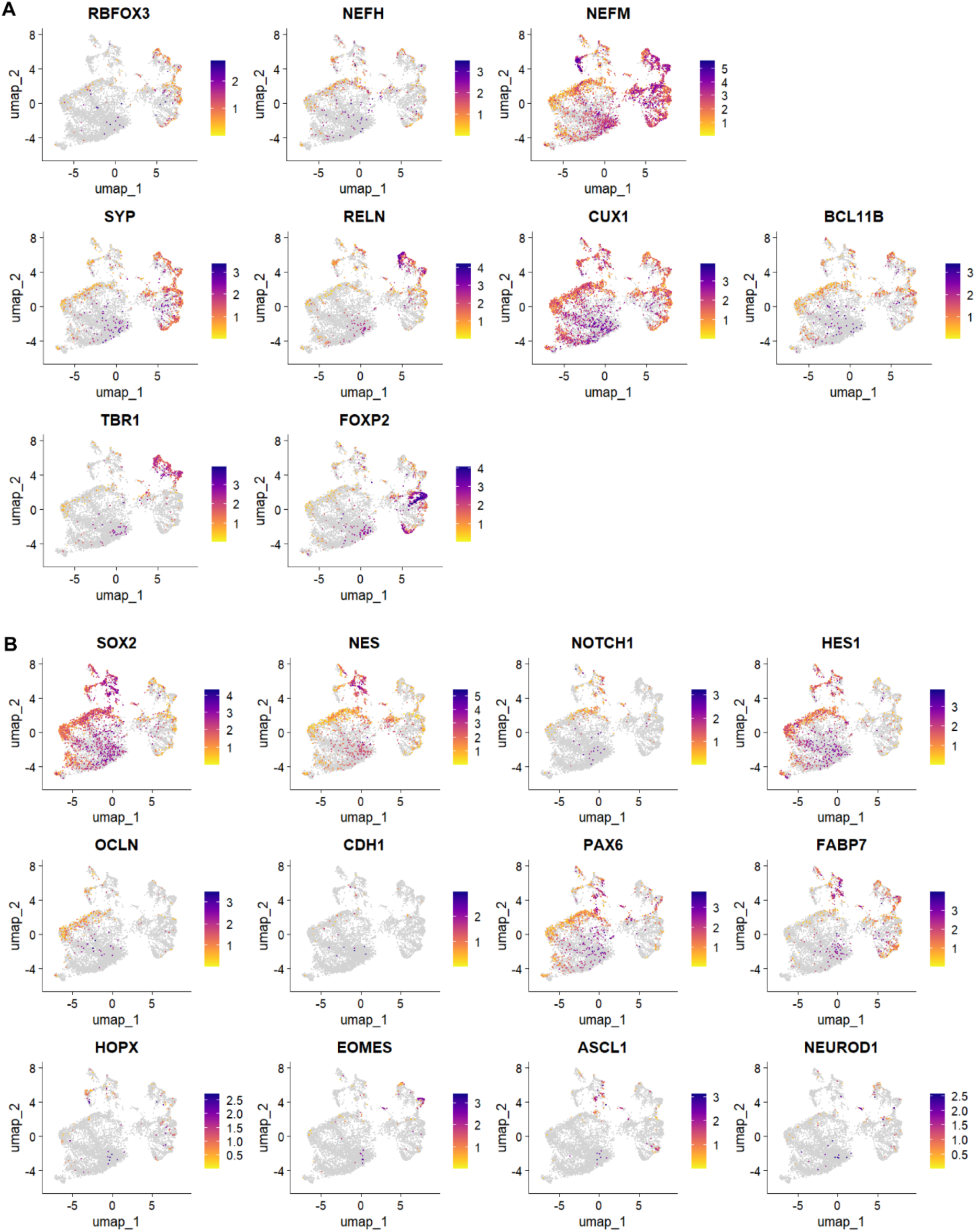
Marker gene expression visualisation from scRNA-seq data. Feature plot of marker genes for progenitor (A) and neuronal (B) marker expression in CPCOs visualised in UMAP. Colour scale represents log transformed counts across cells.

**Supplementary Figure 4.**
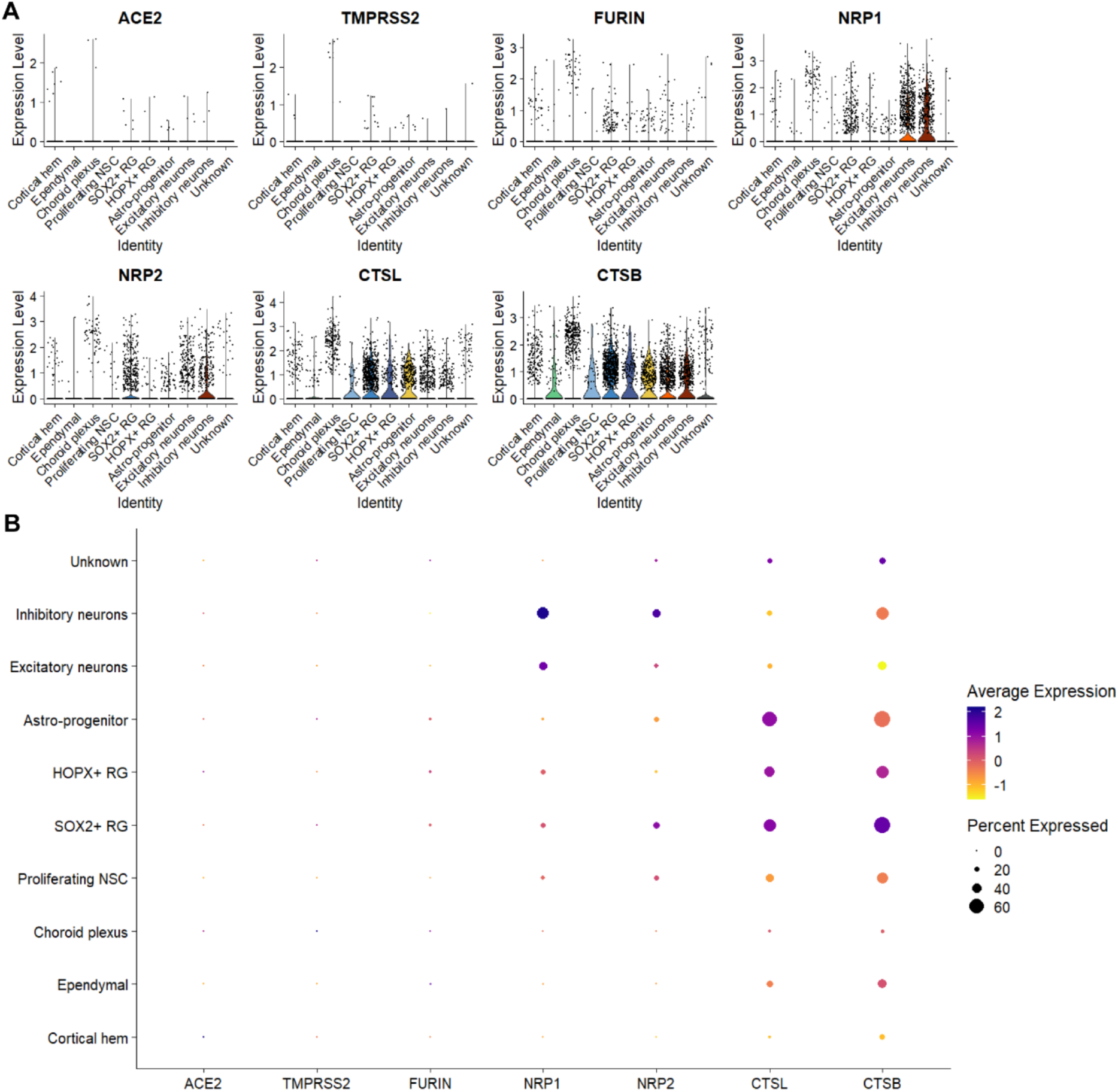
Marker gene expression visualisation from scRNA-seq data. (A) SARS-CoV-2 receptor and protease marker expression profile in CPCOs showed across cell types using violin plot. (B) Selected marker expressions in CPCOs were used to assign cell types across different cell clusters using dot plot, where dots were sized by the percentage of expression and coloured by normalised average expression.

**Supplementary Figure 5.**
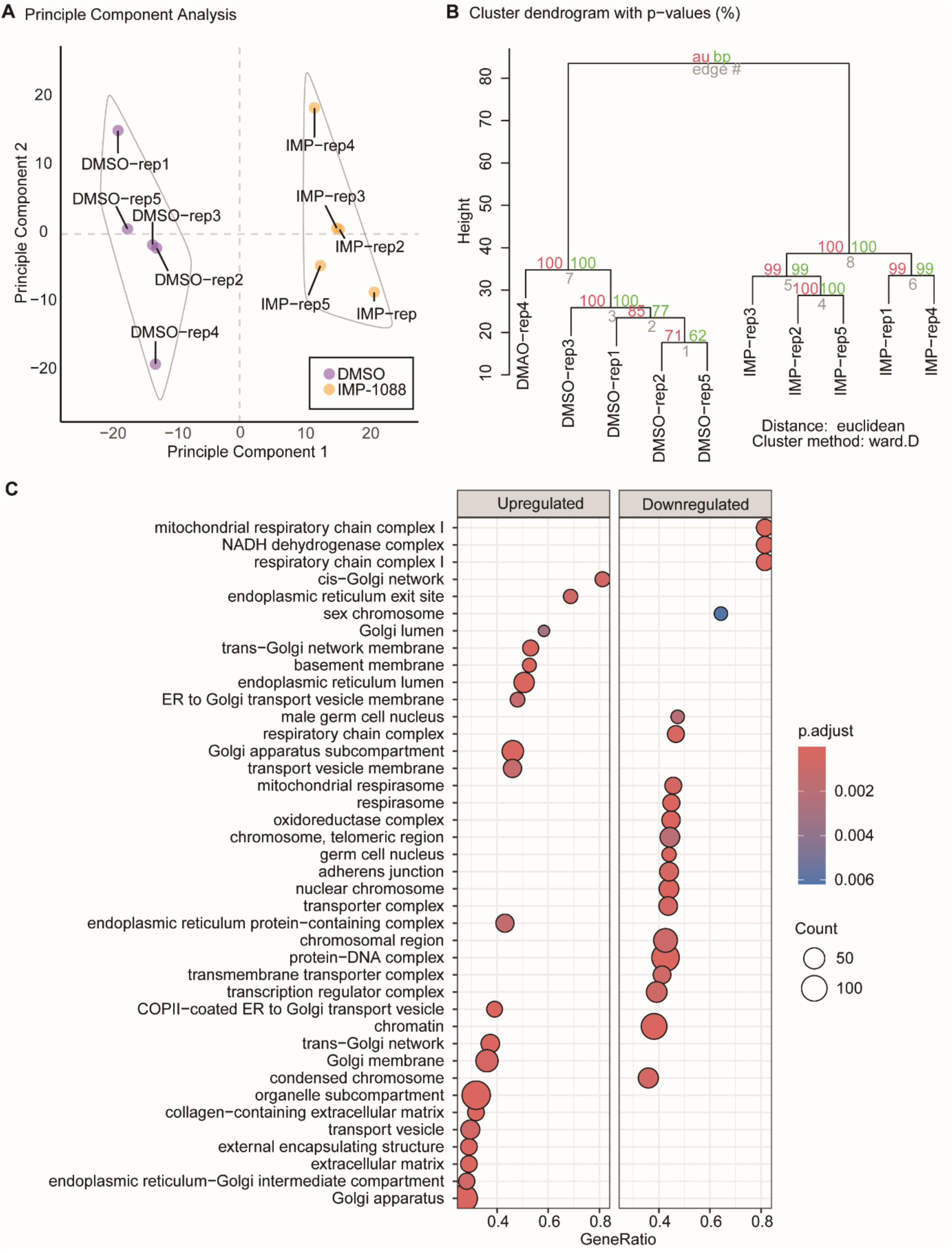
Mass spectrometry analysis of A549-AT cells treated with 1 µM IMP-1088 for 48 h. (A) Principal component analysis. The first two principal components that explained the most amount of variation is shown (x and y-axis). Samples can be observed to cluster according to their experimental group. (B) Hierarchical clustering. Samples cluster according to their experimental group (within each of the tree-like-dendrograms). Values on branches and the red box, represent probabilities of observing the clustering after 100 bootstraps. N = 5 biological replicates in each condition. (C) Over-representation analysis (ORA) for proteins decreased in A549-ATcells treated with 1 µM of IMP-1088 or DMSO control for 48 h revealed Gene Ontology (GO) shows subcellular components on Y axis and gene ratio of upregulated and down regulated protein on X axis with IMP-1088 treatment (DA adjust P<=0.05).

**Supplemental Figure S6:**
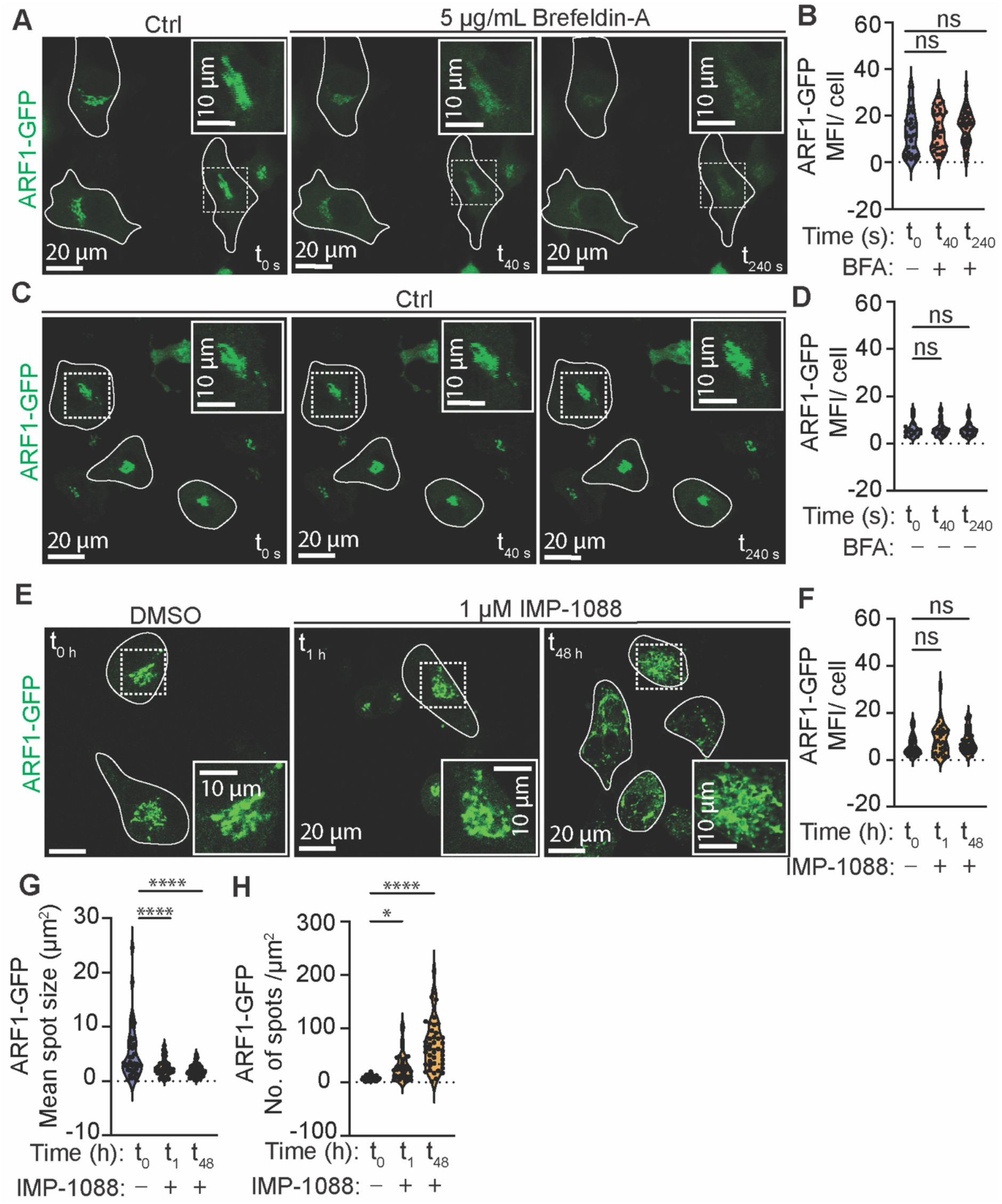
The effects of BFA and IMP-1088 treatments on subcellular localization and distribution of ARF1 are different. (A) Representative live-cell imaging time series of A549-AT cells transiently expressing ARF1-GFP at 0 s (t_0_) and following a treatment with 5 µg/mL Brefeldin-A (BFA) BFA at 40 s (t_40_) and 24 s (t_240_) time points. Boxed areas are enlarged in the corners. Examples of cells are outlined for clarity. (B) Quantification of mean fluorescence intensity (MFI) of ARF1-GFP in A549-AT cells at 0 s (t_0_) and following a treatment with BFA for 40 s (t_40_) and 24 s (t_240_). (C) Representative live-cell imaging time series of A549-AT cells transiently expressing ARF1-GFP at 0 s (t_0_), 40 s (t_40_) and 24 s (t_240_) time points. Boxed areas are enlarged in the corners. Examples of cells are outlined for clarity. (D) Quantification of MFI of ARF1-GFP in A549-AT cells at 0 s (t_0_), 40 s (t_40_) and 24 s (t_240_) timepoints. (E) Representative live-cell images of A549-AT cells transiently expressing ARF1-GFP and treated with vehicle control (DMSO, t_0_) or 1 µM IMP-1088 for 1 h (t_1_) or 48 h (t_48_). Boxed areas are shown on with higher magnification in the corners. Examples of cells are outlined for clarity. (F) Quantification of MFI of ARF1-GFP in A549-AT cells treated with DMSO (t_0_) or 1 µM IMP-1088 for 1 h (t_1_) or 48 h (t_48_). (G) Quantification of ARF1-GFP average spot size (µm^2^) (E) and number of spots per µm^2^ (F) in A549-AT cells in indicated conditions. Violin plots are median ± quartiles. The results are from 3 biological replicates in each condition. Statistical testing was performed with Ordinary one-way ANOVA multiple comparison test. *p<0.05, ****p<0.0001, ns=non-significant.

